# Novel subthreshold retinal laser treatment with ERG-based thermal dosimetry activates hormetic heat response in pig RPE *in vivo*

**DOI:** 10.1101/2022.11.29.518343

**Authors:** Mooud Amirkavei, Ossi Kaikkonen, Teemu Turunen, Anna Meller, Johanna Åhlgren, Anders Kvanta, Helder André, Ari Koskelainen

## Abstract

Boosting natural mechanisms to retain cellular homeostasis and combat oxidative stress by inducing a hormetic heat shock to retinal pigment epithelium with non-damaging transpupillary laser heating, i.e., with subthreshold laser treatment (SLT), has been suggested as a promising therapeutic target for many retinal diseases, including age-related macular degeneration. However, the therapeutic temperature window for the treatments is narrow and thermal dosimetry is unavailable. Here we introduce an SLT modality where the retinal temperature is monitored with electroretinography (ERG)-based thermal dosimetry and demonstrate its feasibility with anesthetized pigs. In 60-second treatments with 810 nm laser, the ED_50_ peak temperature for visible lesion generation was 48°C and the relative temperature determination error was below 10% from the temperature increase. Heat shock protein expression increased, and autophagy was activated at 44.2 °C and no signs of oxidative stress or apoptosis emerged at 44.2 °C or 46.5 °C. The demonstrated method permits a controlled activation of intracellular chaperones and waste clearance in RPE cell with a clear temperature margin for adverse events. In the clinical setting, ERG-based dosimetry would allow safe and personalized SLTs for retinal diseases currently lacking effective treatments.

## Introduction

Age-related macular degeneration (AMD) is a late-onset neurodegenerative retinal disease with clinical and pathological similarities to Alzheimer’s disease (AD) and Parkinson’s disease, including their link to excessive oxidative stress and accumulation of damaged and toxic protein aggregates (Kaarniranta *et al*., 2011). Dry or nonexudative form of AMD is strongly associated with the degeneration and death of retinal pigment epithelium (RPE) cells (Bhutto and Lutty, 2012). These cells provide the neural retina with nutrients, remove waste materials, and perform essential functions for vision (Sparrow, Hicks and Hamel, 2010) while they are exposed to extreme oxidative stress due to the high metabolic rate and oxygen consumption of the retina, and daily light exposure (Sparrow, Hicks and Hamel, 2010).

To cope with exogenous and endogenous stress factors and maintain protein homeostasis, cells have developed highly regulated stress-response mechanisms. In response to stress, such as increased temperature, the heat shock transcription factor HSF1 induces the expression of heat shock proteins (HSPs), which act as chaperone molecules. HSPs detect and refold unfolded or misfolded proteins and suppress the formation of aggregates, a process known as the heat shock response (HSR) (Anckar and Sistonen, 2011). Additionally, HSPs are closely linked to the ubiquitin-proteasome system (UPS) by recognizing misfolded proteins for degradation (Shiber and Ravid, 2014). Once the capacity of HSPs is exceeded or UPS faces functional limitations, another cytoprotective mechanism, autophagy, plays a crucial role in cellular homeostasis by facilitating lysosomal degradation and recycling cytoplasmic protein aggregates (Penke *et al*., 2018).

Investigations of RPE cells from AMD patients have demonstrated a decreased production of heat shock proteins together with deregulation and reduction of autophagic flux (Viiri *et al*., 2013; Mitter *et al*., 2014). Boosting the natural mechanism to retain cellular homeostasis and combat oxidative stress by inducing a hormetic heat shock to RPE has been suggested as a promising therapeutic target for many retinal diseases including AMD (Sramek, Mackanos, Spitler, L. S. Leung, *et al*., 2011; Brader and Young, 2016; Lavinsky *et al*., 2016). Hormesis refers to the stimulation with a substance or stress that is harmful in high quantities but helpful in low levels, such as heat shock. The hormetic zone for inducing heat shock response with subthreshold laser treatments (SLTs) seems to be narrow, since 50–80 % of a damaging thermal dose is needed to increase HSP70 production in RPE and choroid (Desmettre, Maurage and Mordon, 2001; Sramek, Mackanos, Spitler, L.-S. Leung, *et al*., 2011; Amirkavei *et al*., 2020). Additionally, interpatient variability in ocular opacity and blood circulation, RPE and choroidal pigmentation, and morphology of the treated area are known to affect the laser-induced temperature elevation (Mainster and Reichel, 2000).

In recent years, several SLT modalities have emerged and they have early clinical evidence in the treatment of macular edemas due to diabetic retinopathy, retinal vein occlusion, and central serous chorioretinopathy (Scholz, Altay and Fauser, 2017; Wu *et al*., 2018; Gawęcki, 2019), and also in the treatment of AMD (Söderberg *et al*., 2012; Luttrull *et al*., 2015, 2018, 2020). The current state of the art is to use a fixed laser power for all patients or to determine the power individually after titrating a RPE damaging laser power outside the treated area (Manayath *et al*., 2012; Lavinsky *et al*., 2014). However, used treatment parameters and clinical outcomes vary profoundly between clinical trials and no single SLT modality has shown clear superiority (Vujosevic *et al*., 2015; Wang *et al*., 2017; Chhablani *et al*., 2018). Despite the narrow therapeutic window and the variation in individual physiological properties, thermal dosimetry for retinal laser treatments is currently unavailable.

Electroretinography (ERG), a registration of a field potential generated by retinal neurons, is a promising method for thermal dosimetry of SLTs since ERG signaling kinetics are strongly temperature-dependent (Pitkänen, Kaikkonen and Koskelainen, 2017, 2019; Kaikkonen *et al*., 2021). In ERG registration, the retina is stimulated with light and the resulting signal can be recorded non-invasively using corneal and skin electrodes. When utilizing ERG in temperature determination during SLT, a focal ERG (fERG) signal can be elicited directly from the heated area by focusing a light stimulus to the targeted tissue. Laser-induced increases in the retinal temperature can then be calculated by analyzing the acceleration of fERG response kinetics.

This work introduces a subthreshold laser treatment modality for controlled induction of heat shock in the RPE and demonstrates its feasibility in anesthetized pigs in a situation closely resembling the clinical setting for humans. In the method, fERG responses are recorded before and during laser exposure to determine the laser-induced temperature elevation in real-time and to estimate the laser power required to reach a given target temperature. The method was tested with 18 eyes of 11 pigs using an 810 nm infrared laser with a 60-second laser exposure and 3.4- or 5-mm laser spot diameter for the treatment. The treatment sites were investigated for visible lesions, followed by molecular analyses studying the effects of heat shock in the RPE/choroid and neural retinal cells. The steady-state ED_50_ temperature for lesion generation was 47.7 and 48.0 °C and the relative error in the temperature estimate was 13 and 6.8 % for 3.4- and 5-mm laser spot diameters, respectively. In RPE/choroid cells, the non-damaging heat shock for a temperature above 44 °C resulted in the induction of chaperone-related genes followed by increased expression HSP70 and HSP90 proteins. Our results demonstrate that hormetic heat boosts the expression of fundamental autophagy-associated genes and increases the level of LC3B-II protein while p62 protein level decreases, indicating the activation of autophagy in RPE/choroid cells. Oxidative stress and apoptosis were not observed at 44.2 and 46.5 °C in RPE/choroid cells and in the neural retina.

This study highlights that the therapeutic window for subthreshold heat shock is narrow and requires accurate temperature control during heating. Our findings demonstrate that the laser power required to generate harmful lesions can be accurately predicted with ERG-based thermal dosimetry, and the introduced method allows estimation of the personalized power to induce therapeutic effects of SLTs safely and repeatedly. After translation to humans, the presented treatment could be beneficial in the treatment of diseases such as AMD, where the defense systems of aged cells are insufficient to battle against sustained oxidative stress.

## Methods

### Ethical approval

The use and handling of the animals were in accordance with the Finnish Act on the Protection of Animals Used for Scientific and Educational Purposes (497/2013) and the Government Decree on the Protection of Animals Used for Scientific and Educational Purposes (564/2013) and approved by the Project Authorization Board in Finland (project license: ESAVI-30526-2019).

### Animals

11 male domestic pigs (*Sus scrofa domestica*) were acquired from a local farmer. At least 14 days of acclimatization time was provided for each animal before experiments. Animals were at the age of 12–16 weeks (14.8 ± 1.5, mean ± std) and weighted 28 – 55 kg (44.5 ± 8.0, mean ± std) at the time of the experiment day. No signs of sickness or any other symptoms were evident at the time of induction of anesthesia.

Animals were housed in groups at the large animal facility of the Laboratory Animal Centre, Helsinki Institute of Life Science of the University of Helsinki. Toys, straw, and fresh hay were available as enrichments. The solid floor of the pen was covered with wood shavings (Pölkky, Pölkky Oy, Finland). Food (Pekoni 1 mure or Kombi Nasu, Suomen rehu, Hankkija Oy, Finland) was provided twice a day, and fresh tap water was available *ad libitum*. Room temperature was 19 ± 1 °C and relative humidity 45 – 60 %. The pen was cleaned once per day. In addition to natural light from windows, lights were on from 7 am to 15 pm.

### A device for temperature-controlled SLT

The system used for delivering the treatments comprises an ERG-amplifier (FE232 Dual Bio Amp and PowerLab 8/35, 8 Channel Recorder, AD Instruments Ltd., Sydney, Australia), a digitizer and signal generator system (cDAQ-9179, NI-9263 and NI9215, National Instruments, Texas, USA), a laptop computer, a digital slit lamp biomicroscope system (KSL-H5-DR, Keeler, Windsor, United Kingdom), and a fully customized beam generator module mounted on the slit lamp biomicroscope. Additionally, the experimental configuration included a fundus lens (HR wide field, Volk, Mentor, Ohio, USA), mount for the fundus lens (Steady mount, Volk, Mentor, Ohio, USA), a DTL fiber electrode (Unimed Electrode Supplies Ltd, Surrey, United Kingdom), and Ag/AgCl cup electrodes (Reusable EEG cup electrode Silver/Silver Chloride, Technomed, Maastricht, The Netherlands) for reference and ground. The raw ERG signal was amplified and bandpass filtered between 1 Hz and 1 kHz by the ERG amplifier and sampled at 10 kHz.

Fig. 1 A shows a schematic of the beam generator module of the treatment device. The device delivers four independently controlled beams of light onto the fundus through a fundus lens. The four beams are termed the treatment laser beam, the stimulus beam, the central background beam, and the peripheral background beam. The purpose of the treatment laser beam is to elevate the temperature of the target area. The stimulus beam elicits the ERG signal from the target area. The central background beam provides illumination required by fundus imaging, as well as keeps the target area at a steady light adaptation state. The peripheral background beam suppresses ERG signaling elicited by scattered stimulus light in the peripheral retina.

**Figure 1.**
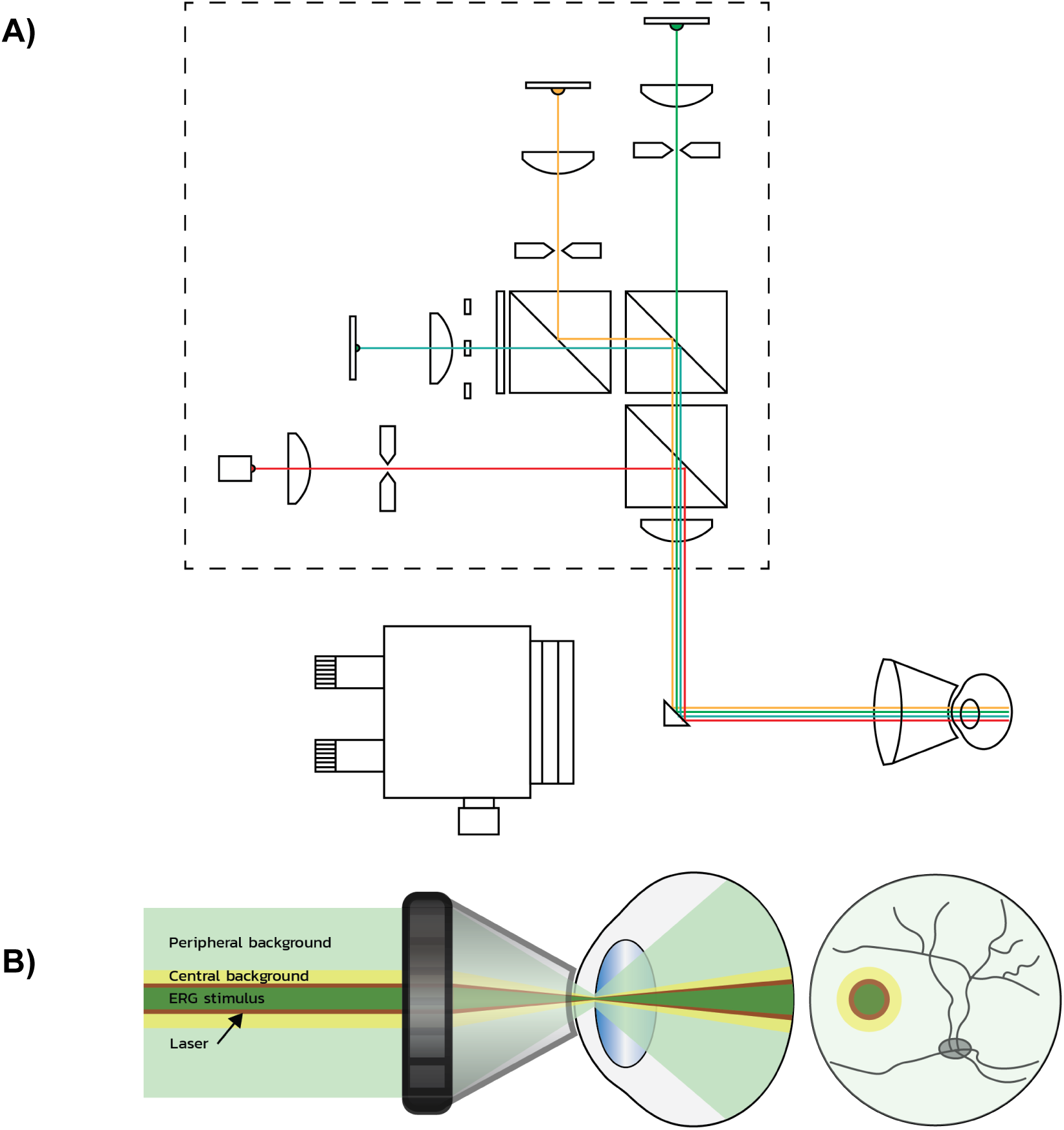
Device schematic. **A)** Beam generator module of the treatment device. The device has four light sources: the laser beam, central background beam, stimulus beam and peripheral background beam. The laser, central background and stimulus beams are first directed to an aperture controlling the spot size, and the aperture is projected to the fundus using a fundus lens. An ophthalmic biomicroscope with a camera is used for fundus imaging. The spectrum of the peripheral background beam is made narrow with an optical bandpass filter and a corresponding band is rejected from imaging with an optical notch filter. **B)** Illustration of beams entering the eye. The central background beam illuminates the target area and provides stable light adaptation, the ERG stimulus beam elicits the ERG signal, the laser beam elevates retinal temperature, and the peripheral background beam light adapts the peripheral retina, suppressing ERG signaling from scattered stimulus light.

Treatments were delivered with laser spot diameters of 3.4 and 5 mm and stimulus spot diameters of 2.7 and 4 mm, respectively. A smaller stimulus spot diameter was used relative to the treatment laser spot to elicit the ERG signal at the center of the laser spot with the highest temperature elevation. The central background beam had a diameter of 8 mm at the fundus, coaxial to the treatment and stimulus beams. The peripheral background was used to illuminate the entire peripheral retina that could be illuminated with the wide-field fundus lens (reported as 160° field of view for humans), except for a unilluminated area corresponding to the treatment laser spot. Fig. 1 B shows the arrangement of different beams on the fundus.

### Light calibrations

The porcine cone system consists of two types of cone cells; medium (M) and short (S) wavelength cones with maximal absorption at 556 and 439 nm, respectively (Neitz and Jacobs, 1989). The S cones represent a minority, between 7.4 and 17.4 % of all cone cells (Hendrickson and Hicks, 2002). The white central background beam light adapts both the S and M cones, driving their sensitivity down, while the stimulus light excites primarily the M cones. Due to the small proportion of S cones and the selection of stimulus and background wavelengths, the ERG signal originates nearly exclusively from M cones. Hence the photometric units used in this study were calibrated for the porcine M cone by using the sensitivity spectrum template devised by (Govardovskii *et al*., 2000).

The beam irradiances were measured by placing an optical power sensor (PM121D, Thorlabs, Newton, NJ, USA) at the focal plane prior to the fundus lens, while homogeneity of the beams was verified using a beam camera (Laser Beam Diagnostics SP503U, Spiricon). The measured irradiance was corrected using the sensitivity spectrum of the sensor and the spectra of the light sources and converted to illuminance using the sensitivity spectrum of the porcine M cone. The magnification of the Volk HR wide-field fundus lens was determined to be 1.35 for porcine eyes by comparing the spot sizes to the size of the optic nerve during the treatment procedure and determining the optic nerve diameter after enucleation. The reported retinal illuminance and irradiance, as well as spot diameter values, take this magnification into account.

### Light stimulation and ERG signal processing

The retina was stimulated either with a 16 Hz flash flicker or with a flicker with uneven flicker intervals with an average frequency of 17 Hz. The uneven flicker interval comprised 16 different flash intervals evenly spaced between 40 and 75.5 ms and the order of the flash intervals was altered pseudorandomly. The duration of a single flash stimulus was 1 ms. The retinal illuminances of the central and peripheral background beams were kept constant at 1150 and 2300 lx, and the luminous exposure delivered in a single flash stimulus was kept at 46 lx·s during all laser treatments. The retinal illuminance caused by the treatment laser beam was below 50 lx in all laser treatments. As the central background beam provides a > 20x brighter background, light adaptation caused by the laser beam was deemed insignificant.

As fERG responses to single flash stimuli are dominated by noise, a signal containing a series of responses has to be processed to produce a fERG response with a sufficient signal-to-noise ratio (SNR) for retinal temperature determination. A denoised fERG signal f(n) was computed using the cross correlation between the recorded fERG signal y(n) and a stimulation vector s(n), shown in Eq. 1. The stimulation vector contains ones where the flash stimulus was fired and zeros elsewhere. This method produces identical results to signal averaging when computing the denoised ERG response using a constant stimulation frequency, but uneven flicker intervals allow computing the full-length impulse response.

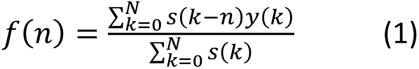

### ERG-based thermal dosimetry

#### Retinal temperature determination

The temperature determination algorithm, shown in Eq. 2 and described with further detail in (Kaikkonen *et al*., 2021), takes two ERG responses *f*_1_(*t*) and *f*_2_(*t*) as inputs, with t = 0 set to the moment of the flash stimulus, and determines the difference in retinal temperature Δ*T* between two ERG responses. A value of 3.7 %/ °C was used for the temperature dependence of kinetics acceleration, as determined for pig cones with similar stimulus-background illumination contrast in (Kaikkonen *et al*., 2021, their Fig. 5b). Similar temperature dependence has been demonstrated also for mouse scotopic ERG signal kinetics (Pitkänen, Kaikkonen and Koskelainen, 2017, 2019)

The algorithm compresses or expands the time axis of one of the responses an amount that maximizes the Pearson correlation between the compared responses and uses this value as a temperature-dependent feature. The difference in retinal temperature between the responses is proportional to the logarithm of this feature.

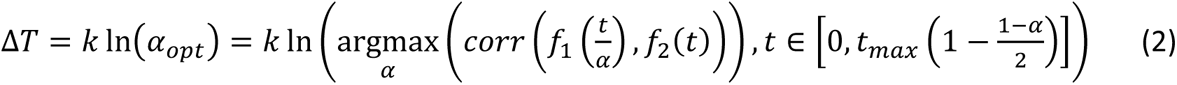

For each laser exposure the laser-induced temperature elevation was determined by comparing an ERG response acquired during laser exposure, henceforth termed exposure ERG response, to an ERG response recorded directly before the laser exposure when the retina is at body temperature, termed pre-exposure reference ERG response. The ERG responses acquired after the first 15 seconds of laser exposure were used to compute the denoised ERG response for temperature determination to reflect the steady-state temperature elevation induced by the laser exposure.

#### ERG-based laser power calibration

The local thermal responsivity, i.e., the amount of laser power required per unit temperature increase, was determined before each treatment through a calibration protocol. The calibration protocol involved delivering three laser exposures with different powers causing < 5 °C increases in retinal temperature. The retinal temperature increase caused by each of the exposures was determined and a zero-intersecting line was fitted between the laser powers and the determined temperature elevations. The local thermal responsivity was defined as the slope of the fitted line. This thermal responsivity value was used to determine a laser power that leads to a preset target temperature elevation, and a 60-second laser exposure was performed with the determined laser power.

Two different calibration protocols were used in this study. In the first protocol, the retinal temperature was elevated with three separate 60-second laser exposures, allowing the retinal temperature to return to body temperature before the subsequent laser exposure (Fig. 2 A). In the second calibration protocol, the pre-exposure reference ERG response was acquired first, followed by three 30-second laser exposures with different powers without turning the laser off in between (Fig. 2 B). In both protocols, the temperature elevations caused by the calibration laser exposures were calculated from the acceleration of the ERG responses.

**Figure 2.**
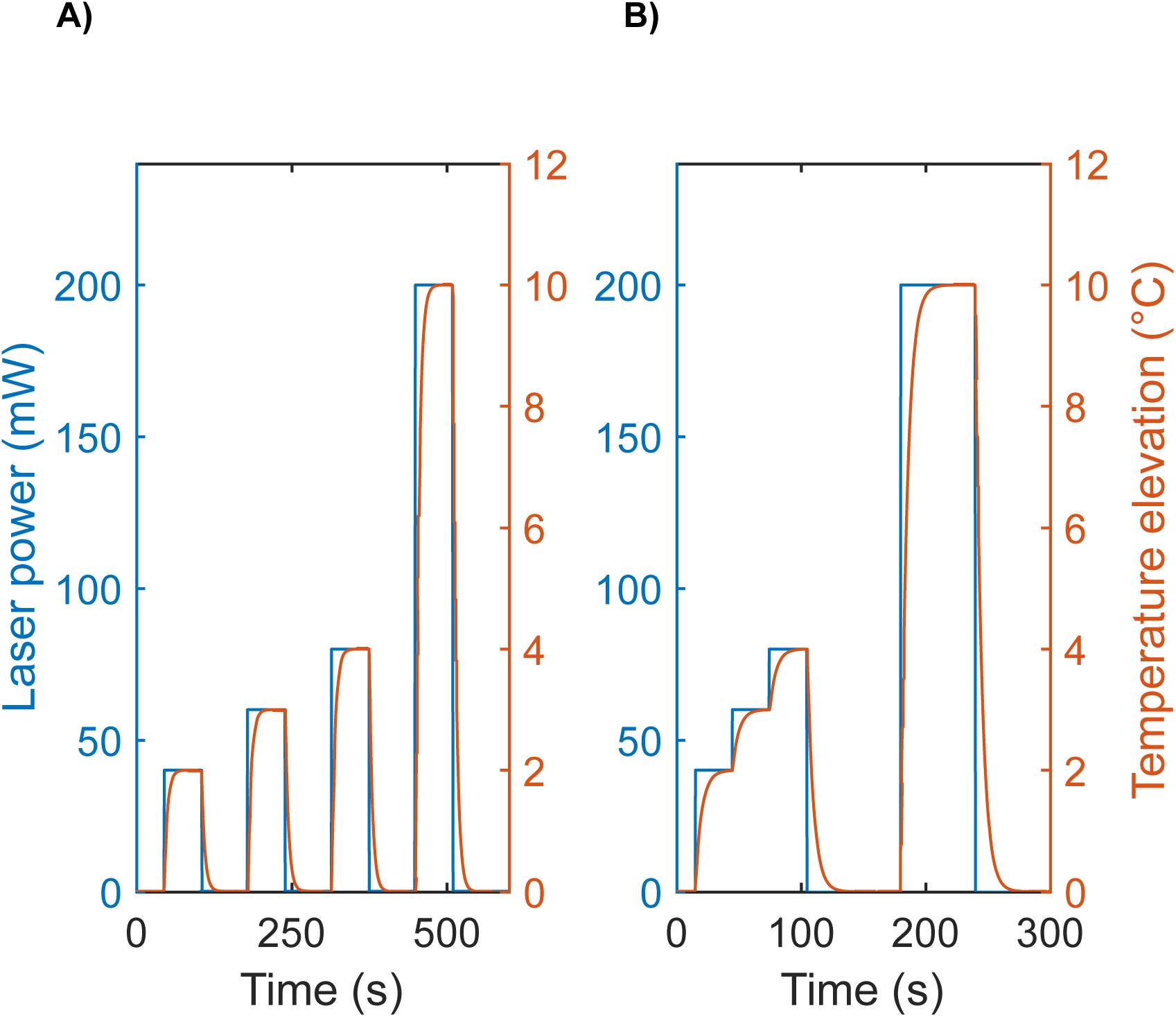
Illustration of the individualized laser power calibration protocols. **A)** The protocol comprises three separate 60-second laser exposures, allowing retinal temperature to normalize in between. fERG responses recorded before laser application are compared to fERG responses recorded after the first 15 seconds of laser exposure to determine the steady state temperature increase. B) The protocol comprises three consecutive 30-second laser exposures. fERG responses recorded before the first calibration exposure are compared to fERG responses recorded at the last 15 seconds of each consecutive laser exposures. The calibration protocols are followed by 60-second laser treatment.

### Experimental procedures

#### Pig anesthesia

The anesthesia was induced as described in (Kaikkonen *et al*., 2021). Before each anesthesia, the pigs fasted at least 17 hours. The anesthesia was maintained by total intravenous (IV) anesthesia (TIVA), using constant rate infusions (CRI) of propofol (2.0 – 4.8 mg/kg/h), fentanyl (0.003 – 0.006 mg/kg/h) and midazolam (0.3 – 0.7 mg/kg/h). The infusion rates were adjusted to meet the individual needs of each animal by constantly monitoring the anesthesia level and the vital parameters (heart rate, respiratory rate, saturation, end tidal CO_2_).

The first day anesthesia lasted from 4 hours 15 minutes to 6 hours (from intubation to extubation). The pigs were placed on a prone position on the table. The eyes of the anesthetized pig were moistened repeatedly during the experiments with Viscotears 2 mg/g eye gel (Dr. Gerhard Mann chem.-pharm. Fabrik GmbH, Berlin, Germany). The eye that was not currently treated was closed with tape. The pupils were dilated with tropicamide (Oftan® Tropicamide 5 mg/ml, Santen Oy, Tampere, Finland) and the corneas were anesthetized with oxybuprocaine hydrochloride (Oftan® Obucain 4 mg/ml, Santen Oy, Tampere, Finland), respectively. With few individuals the eye shifted from normal position. The eye position was reversed to normal by deepening the anesthesia level either by increasing the flow rates or giving boluses of ketamine. (1-2 mg/kg iv)

After the treatment procedures, the pigs were gradually weaned from the ventilator and extubated when breathing spontaneously. The need for reversal drugs was evaluated individually. If needed, atipamezole (0,2 mg /kg) and flumazenil (0,003-0,007 mg/kg) were given. The pigs were moved to recovery area, oxygenated, and monitored until full recovery.

Pigs were let to follow their normal awake/sleep rhythm for 18 – 20 hours before the second anesthesia, which followed the same procedure as the anesthesia during the first day. At the end of the procedure, the pigs were euthanized by 100 mg/kg pentobarbital IV. The anesthesia at the second day lasted from 1 hour 40 minutes to 5 hours (from intubation to sacrificing).

#### Preparations

##### Temperature monitoring

A nasopharyngeal temperature probe, inserted ca. 10 cm deep into the nasal cavity (DMQ-DAG-20-N0, Cables and Sensors, Orlando, USA), was used for continuous monitoring of body temperature. Fluctuations of nasopharyngeal temperature was minimized by blocking the gas exchange through nostrils. The body temperature was maintained constant (38.5 ± 0.6 °C, mean ± std) using Bair Hugger™ patient warming system (3M, Minnesota, United States) where an air mattress (Bair Hugger™ Pediatric Full Body Blanket, 31000), with a constant flow of warmed air through it, was placed on the pig.

##### ERG electrode placement

The places for the ERG reference and ground electrodes were shaved, cleaned, and scrubbed with abrasive gel (Nuprep, Weaver and Company, Colorado, USA) to improve the skin conductivity. The reference electrode was attached to the corner of the eye and the ground electrode to the pig ear with conductive paste (Ten20, Weaver and Company) and tape. DTL thread electrode was inserted on the eye at the rim of the lower eye lid.

#### Laser treatments

After the preparations, a contact fundus lens was placed on the eye and the biomicroscope was focused on the fundus, which also focuses the treatment and stimulus beams. A retinal area was selected, and the calibration protocol was performed to determine the laser power required for unit temperature elevation at the selected area. The laser power for the treatment was chosen based on the calibration protocol to reach a target temperature set for the treatment. The length of the treatment was always 60 seconds, and ERG responses were continuously monitored during the treatment. The treatment location and possible eye movements were followed from the live feed fundus view from the camera connected to the biomicroscope.

##### Treatment set 1 – Determination of therapeutic temperature window

Total of 36 laser treatments were first conducted with 10 eyes of 7 pigs to determine the therapeutic temperature window for HSP70 production and generation of lesions. The target temperatures ranged from 41 to 51 °C. The diameter of the laser spot was 3.4 mm and of the stimulation spot 2.7 mm at the fundus. Constant 16Hz flicker and uneven flicker light stimulation protocols were used in 26 and 10 treatment spots, respectively. The calibration protocol to determine the thermal responsivity of the target area comprised three separate 60-second laser exposures, allowing the retinal temperature to return to body temperature after each exposure (see Fig. 2 A). During heating, the first 15 s were reserved for temperature stabilization and the recording duration for the pre-exposure reference and exposure ERG responses was 45 seconds in both the calibration and treatment protocols.

##### Treatment set 2 – Quantitative analysis of heat-induced biochemical effects

After the first set of experiments, the laser and stimulus spot diameters were increased to 5 mm and 4 mm, respectively, to increase the signal-to-noise ratio of ERG recording and to produce enough material for quantitative molecular analysis from individual treated spots. Target temperatures were divided to three groups having the mean temperatures of 44.2, 46.5 and 48 °C. The set comprised 36 treatment areas in 8 eyes of 4 pigs. Uneven flicker ERG stimulation protocol was used in all treatments. Thermal responsivity of the target area was calibrated by performing three consecutively 30-second laser exposures without allowing retinal temperature to lower in between (see Fig. 2 B). The first 15 seconds were reserved for temperature stabilization after laser power changes, and the recording duration of the pre-exposure reference and exposure ERG responses was 15 seconds for both the calibration and treatment protocols.

#### Anterior eye damage and general habitus of animals

After completing all treatments on both eyes, I-DEW FLO Fluorescein Strips with NaCl 0.9 % were used to diagnose possible abrasions or wounds on the cornea. The animals were allowed to emerge from anesthesia while being monitored until the animal was standing and eating. General habitus and irritation in eyes were monitored again in the morning of the following day. Irritation was diagnosed in the case of redness, scratching, or squinting of the eye.

#### Fundus imaging and Sample preparations

The cornea and the iris of both eyes were photographed before pupil dilation with a corneal imaging module (ES2 eye surface lens, Smartscope VET, Optomed, Oulu, Finland) connected to a handheld fundus camera (Smartscope Pro, Optomed, Oulu, Finland) and color fundus images were taken after pupil dilation using the same camera and retinal module (EY4, Smartscope VET, Optomed, Oulu, Finland) at the beginning of the first and second day anesthesia to examine any treatment-induced damage. Additionally, fluorescence angiographic (FA) images were taken on the second day with a FA module connected to the fundus camera (FA lens, Smartscope VET, Optomed, Oulu, Finland) after injection of 2.5 ml of fluorescein sodium (Fluoresceine 10% Faure, SERB SA, Paris, France) to the cannulated auricular vein. The injection of 2.5 ml of fluorescein was repeated before taking the FA images from the second eye. After photographing, the pig was euthanized, and the eyes were isolated and stored to a cold PBS in a cooler filled with ice. The eyes were measured and opened within 30 minutes from euthanasia, and images were taken from the open eye cup with a digital camera (Nikon D800, Nikon, Japan, and Sigma 105 mm 1:2.8 DG Macro HSM, Sigma, Japan).

In treatment set 1, after enucleation of the eye, the posterior eye cup containing the RPE/choroid/sclera were isolated for immunofluorescence staining (see next chapter for details). In treatment set 2, the locations of the laser treatments were punched with 5 mm biopsy needle from the eye cup, and the gained neural retina and RPE/choroid samples were stored separately in liquid nitrogen. Additionally, 4 – 6 non-treated samples were collected from each eye and considered as non-treated controls for the molecular analysis.

### Immunofluorescence

Immunofluorescence staining of the posterior eyecup containing RPE, choroid, and sclera was conducted in order to evaluate the HSP70 expression at different temperatures. Eye cup was rinsed with PBS, fixed with 4 % formaldehyde (FA; Solveco, Rosersberg, Sweden) in phosphate-buffered saline (PBS; Gibco, Paisley, UK) for 30 min at room temperature (RT), following by permeabilization with 0.1 % Triton X-100 (Sigma-Aldrich Corp.) in PBS for 30 min at RT. Permeabilized eye cups were incubated in a blocking solution of 10 % normal goat serum (Invitrogen, Camarillo, MD, USA) supplemented with 0.1 % Triton X-100 for 1 h at RT. Then, the samples were incubated with mouse primary antibody HSP70 (1:50, cat. no SPA-810; Enzo Life Science, Santa Cruz Biotechnology, Paso Robles, CA, USA) diluted in blocking solution overnight (ON) at 4 °C. Incubation with secondary antibodies diluted in blocking solution was performed for 2 h at RT: Anti-mouse-Alexa 647 (1:500, cat. A32787, ThermoFisher Scientific Inc.), Alexa Fluor 488-Phalloidin (1:500, cat. no A12379, ThermoFisher Scientific Inc.), Hoechst 33258 (1:10000, Sigma-Aldrich Corp.). Eye cups were extensively washed with PBS after each antibody steps. Finally, eye cups were post-fixed for 10 min in 4 % FA-PBS at RT following by flat-mounting with fluorescent mounting medium (Dako, Carpinteria, CA, USA). Fluorescence images were acquired with Nikon Eclipse Ti-E microscope (Nikon, USA).

### RPE Cell Viability: Tunel Assay

To visualize the apoptosis in RPE cells, a terminal deoxynucleotidyl transferase-mediated dUTP nick-end labeling (Tunel) imaging assay kit (Clickit Plus Tunel 594, Thermo Fisher, USA) was used according to the manufacturer’s protocol. Eye cups were prepared as such for immunofluorescence. Images of the RPE side of the eyecup flat mounts were acquired with Nikon Eclipse Ti-E microscope (Nikon, USA).

### Reactive oxygen species (ROS) quantification

The level of reactive oxygen species was assessed in the homogenate supernatant of RPE and retina tissue with using 2′–7′-dichlorofluorescein diacetate (DCFH-DA; ThermoFisher Scientific Inc.) molecular probes. Briefly, an aliquot of the sample containing 8 µg protein was incubated with 10 µM DCFH-DA for 40 min at 37 °C; the reaction was terminated by cooling the mixture in ice. Measurement of autofluorescence for the sample and reagent was processed with using either an aliquot of sample without DCFH-DA or DCFH-DA without sample in the same experimental condition. In this assay, acetate group of DCFH-DA are removed by esterases and produces the reduced form DCFH, which can be oxidized by ROS to fluorescent oxidized DCF. Formation of the oxidized fluorescent derivative (DCF) was immediately measured with a fluorescence spectrophotometer (TECAN, F-200) at excitation and emission wavelengths of 488 and 525 nm, respectively. All procedures were conducted in darkness. Non-treated tissue served as control. The results were presented as percentage of DCF fluorescent expression relative to control.

### Quantitative reverse transcriptase PCR (qPCR) and data analysis

Total RNA was extracted and purified from RPE tissue punches using Single Shot Cell Lysis Kit (Bio-Rad Laboratories). cDNA synthesis was performed from 1.5 μg RNA using RT² First Strand Kits (Qiagen, Hilden, Germany). Custom RT² Profiler PCR Arrays and SYBR® Green PCR Master Mix (Qiagen, Hilden, Germany) were used to detect expression of transcripts. Each array was designed to profile the expression of 91 transcripts (for a complete list of analyzed genes see Supplementary Table 1). qPCR was conducted using CFX real-time thermal cycler system (Bio-Rad Laboratories). Cycle threshold (Ct) of each replicate was normalized to the average Ct of 2 housekeeping genes: Glyceraldehyde 3-phosphate dehydrogenase (GAPDH) and (β)-actin (ACTB) on a per plate basis. Relative transcript expression for treated samples were normalized to non-treated controls (NTC) applying the ΔΔCT method from 10, 10 and 12 replicates for NTC, 44.2 °C and 46.5 °C, respectively.

### Immunoblotting

To conduct immunoblotting, punches of RPE and retina tissue from the location of treatments were isolated. Tissue punches were suspended in protein extraction RIPA lysis buffer (Sigma-Aldrich Corp) containing protease and phosphatase inhibitors cocktail (Roche, Mannheim, Germany) for 10 min on ice. Tissues were homogenized by 3 × 10 s pulses at maximum speed with a VDI12 rotorstator (VWR, Dresden, Germany). Protein extracts were quantified by a Bradford protein assay (Bio-Rad Laboratories, Hercules, CA, USA). Fifteen micrograms total protein extracts were separated by 4–20 % stain-Free SDS-PAGE gels and transferred onto polyvinylidene difluoride (PVDF) membranes (Bio-Rad Laboratories). Membranes were blocked using Tris-buffered saline (TBS) with 1 % casein (Bio-Rad Laboratories) for 1 h at room temperature. Subsequently, membranes were incubated ON at 4 °C with primary antibodies diluted in blocking solution supplemented in 0.05 % Tween-20 (Sigma-Aldrich Corp.); anti-HSP70 (1:500, mouse polyclonal, cat. no SPA-810; Enzo Life Science, Santa Cruz Biotechnology, Paso Robles, CA, USA); anti-p62 (1:500, rabbit monoclonal, cat. no. NBP1-48320; Novus Biologicals, Centennial, Colorado, USA); anti-LC3B (1:250, rabbit monoclonal, cat. no. 3868; Cell Signaling Technology, Danvers, MA, USA), anti-HSF1 (1:500, rabbit monoclonal, cat. no. 12972; Cell Signaling Technology, Danvers, MA, USA), anti-caspase 8 (1:250, rabbit polyclonal, cat. no. ab227430; Abcam, Cambridge, UK), anti-HSF2(1:250, rat monoclonal, cat. no. NB110-96434; Novus Biologicals, Centennial, Colorado, USA), anti-HSP90 (1:500, rat monoclonal, cat. no. SPA-835; Enzo Life Science, Santa Cruz Biotechnology, Paso Robles, CA, USA), anti-VEGF165 (1:500, mouse monoclonal; Sigma-Aldrich Corp.). Incubation with secondary antibodies diluted in blocking solution was performed 1 h at RT: anti-rabbit-IgG conjugated to horseradish peroxidase (1:10,000, cat. no. P044801-2; Dako, Carpinteria, CA, USA), anti-mouse-IgG conjugated to horseradish peroxidase (1:10,000, cat. no. P016102-2; Dako, Carpinteria, CA, USA), anti-rabbit StarBright 700 (1:2500, cat.no. 12004161, Bio-Rad Laboratories, Hercules, CA, United States), anti-mouse StarBright 520 (1:2500, cat.no. 12005866, Bio-Rad Laboratories, Hercules), hFAB Rhodamine anti-Actin (1:2000, cat.no. 12004164, Bio-Rad Laboratories, Hercules, CA, United States), anti-β-actin (1:5000, rabbit monoclonal, cat. no. SAB5600204; Sigma-Aldrich). Membranes were extensively washed with TBS-T (TBS supplemented with 0.05 % Tween-20) after all antibody steps. Finally, the protein of interest was visualized either by enhanced chemiluminescence (Clarity ECL; Bio-Rad Laboratories) or direct fluorescence with a Chemidoc MP imaging system (Bio-Rad Laboratories). Image Lab 3.0 software (Bio-Rad Laboratories) was used to determine the optical density (OD) of the bands and protein levels correction was against actin as loading control.

### Data and Statistical Analysis

#### Probit model

The retinal temperature of each of the treatments, *T*_*n*_, was estimated with the ERG-based laser power calibration protocol. The temperature estimate *T*_*ERG*,*n*_ is expected to contain a normally distributed error with standard deviation *σ*_*T*_.

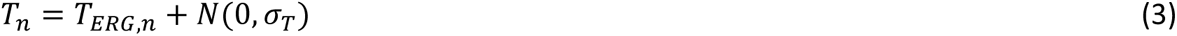

The formation of a visible lesion was modeled as a binary event that occurs above a certain threshold temperature for the treated area *T*_*lesion*,*n*_, which is expected to be normally distributed around the ED50 temperature for lesion generation.

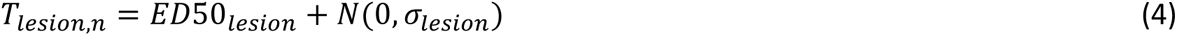

Each treatment was labeled as either causing or not causing a lesion (*L*_*n*_ = 1 for lesion, *L*_*n*_ = 0 for no lesion). Hence, the probability for a given treatment to cause a lesion, shown in Eq. 5, corresponds to the likelihood that the actual retinal temperature was above the lesion threshold temperature for the treated area.

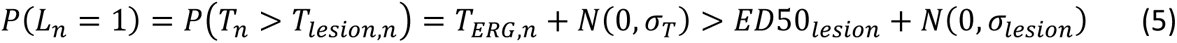

The error terms for retinal temperature determination and possible deviations in the lesion threshold temperature are combined into a single error term 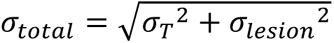. The probability of generating a lesion is shown in Eq. 6 using the cumulative normal distribution Φ with mean *ED*50_*lesion*_ and standard deviation *σ*_*total*_.

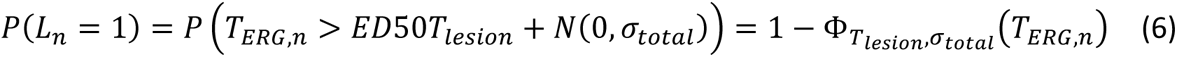

Using a flat prior, the posterior distribution of *ED*50_*lesion*_and *σ*_*total*_ is the same as the likelihood of the observed data.

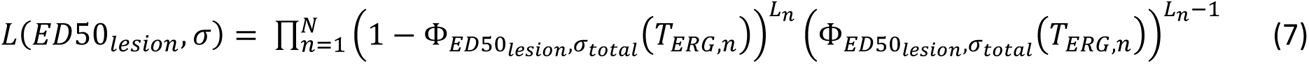

The reported ED50 temperatures for lesion generation and the standard error in the determined retinal temperature are the values that maximize this likelihood function. The *σ*_*total*_ can be seen as an upper bound for the error in retinal temperature determination, as it contains deviations in the lesion threshold temperature as well as the temperature determination error. The reported 95% confidence intervals refer to the intervals of *ED*50_*Lesion*_ and *σ*_*total*_ that correspond to 95% of total likelihood.

The same probit model was also used to determine a threshold temperature for triggering HSP70 production visible in immunostaining. The treatment was considered to increase HSP70 expression if the relative fluorescent intensity increase in the treated area compared to the surrounding areas was 2-times larger than the relative standard deviation of the fluorescent intensity in the surrounding areas in the flat mounted immunofluorescent images (analyzed with Image J 1.52p freeware software, Wayne Rasband, National Institutes of Health, Bethesda, Maryland, USA).

#### Molecular statistical analysis

Molecular statistical analyses were calculated using Prism software (GraphPad Inc., San Diego, CA, USA). All data were subjected to one-way analysis of variance (ANOVA), corrected by Sidak or Bonferroni post-hoc tests for multiple comparisons, as described in figure legends. Data were presented as mean ± standard error of mean (SEM), values of p < 0.05 were considered statistically significant.

## Results

### Focal ERG allows *in vivo* retinal temperature determination during laser-induced hyperthermia

The optimal fERG stimulation for temperature determination should elicit high amplitude responses with a fast signal repetition rate to maximize the signal-to-noise ratio and temporal resolution of the method. Fig. 3 A shows exemplary fERG impulse responses recorded with a 4 mm diameter stimulus spot using the uneven flicker protocol (see Methods). The three responses were elicited with flash stimuli with luminous exposures of 23, 46, and 92 lx·s against a constant white background with 1150 lx illuminance. The white background was delivered in an 8 mm diameter circular spot, which also enabled visualization of the highly pigmented pig fundus during the experiments. Additionally, a peripheral background light of 2300 lx, was projected to the fundus to minimize the effect of stray light to the fERG signal (see Fig. 1 B for optical configuration). A luminous exposure of 46 lx·s was selected for fERG stimulation during laser treatments because it elicits high amplitude non-saturated fERG responses with the used background illumination. The fERG response amplitude per stimulation area, measured from the peak of the a-wave to the peak of the b-wave, was 220 ± 73 nV/mm^-2^ (mean ± std, n = 72) with the chosen stimulus strength at normal body temperature.

**Figure 3.**
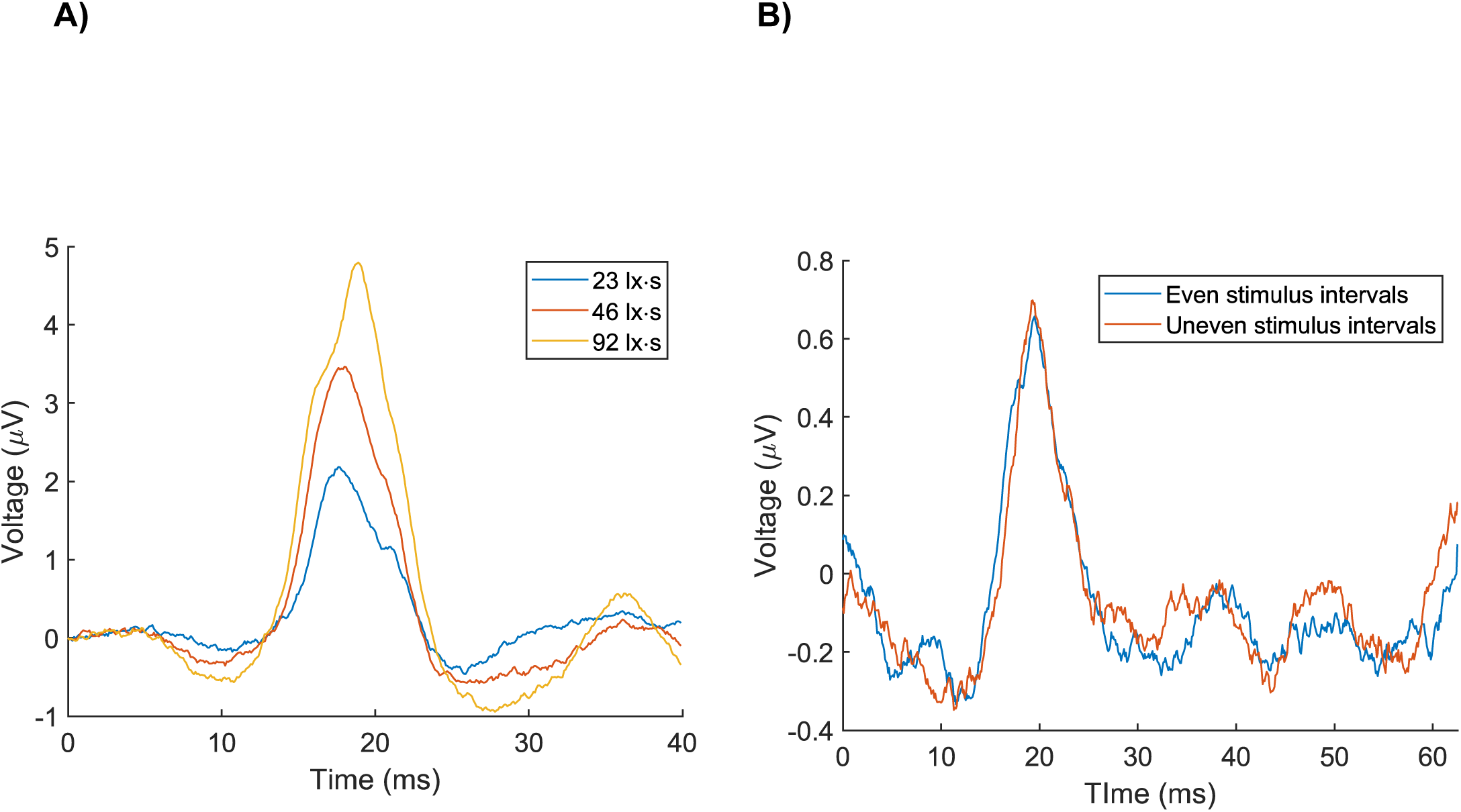
Focal ERG from pig cone photoreceptors. **A)** fERG responses acquired with three flash strengths. **B)** Comparison of the 16 Hz flicker stimulation (blue trace) with uneven flicker protocol (red trace). Signal distortion due to overlapping signals is visible in the early negative wave collected with steady 16 Hz flicker but does not appear in the trace collected using the uneven flicker protocol.

Occasionally, persistent oscillating waveforms tailed the b-wave of the fERG signal and overlapped with the a- and b-waves generated by the subsequent flash. Typically, the effect of signal overlap was mild but, in rare cases, oscillations significantly affected the early fERG waveforms elicited with a steady 16 Hz flickering stimulus. An uneven flicker protocol was deployed instead of a steady flicker to reduce the effect of the oscillations after the experiments with 6 pigs (see detailed description in Methods). Fig. 3 B shows fERG responses from the same 2.7 mm diameter retinal area elicited with a steady 16Hz flicker and with the uneven flicker protocol. The effect of the signal overlap is visible mainly in the early part of the blue trace recorded with steady flicker but disappears when using the uneven flicker protocol (orange trace). Results obtained with uneven flicker, as well as results with steady flicker, where the effect of signal overlap was mild, were not separated in further analysis. However, results obtained with the constant frequency flicker with extremely strong signal overlap were disregarded.

To estimate the temperature increase during laser treatments, fERG signal was continuously recorded from the heated area. The temperature increase was calculated based on the acceleration of fERG response kinetics by comparing the fERG recorded during laser heating to a reference fERG response recorded in the body temperature (see Methods for details) (Pitkänen, Kaikkonen and Koskelainen, 2017, 2019; Kaikkonen *et al*., 2021). Fig. 4 A illustrates fERG responses recorded without laser application and with four different intensities of laser heating applied to the same location. Fig. 4 B shows the steady-state temperature increase estimates with the concerned laser exposures (black crosses).

**Figure 4.**
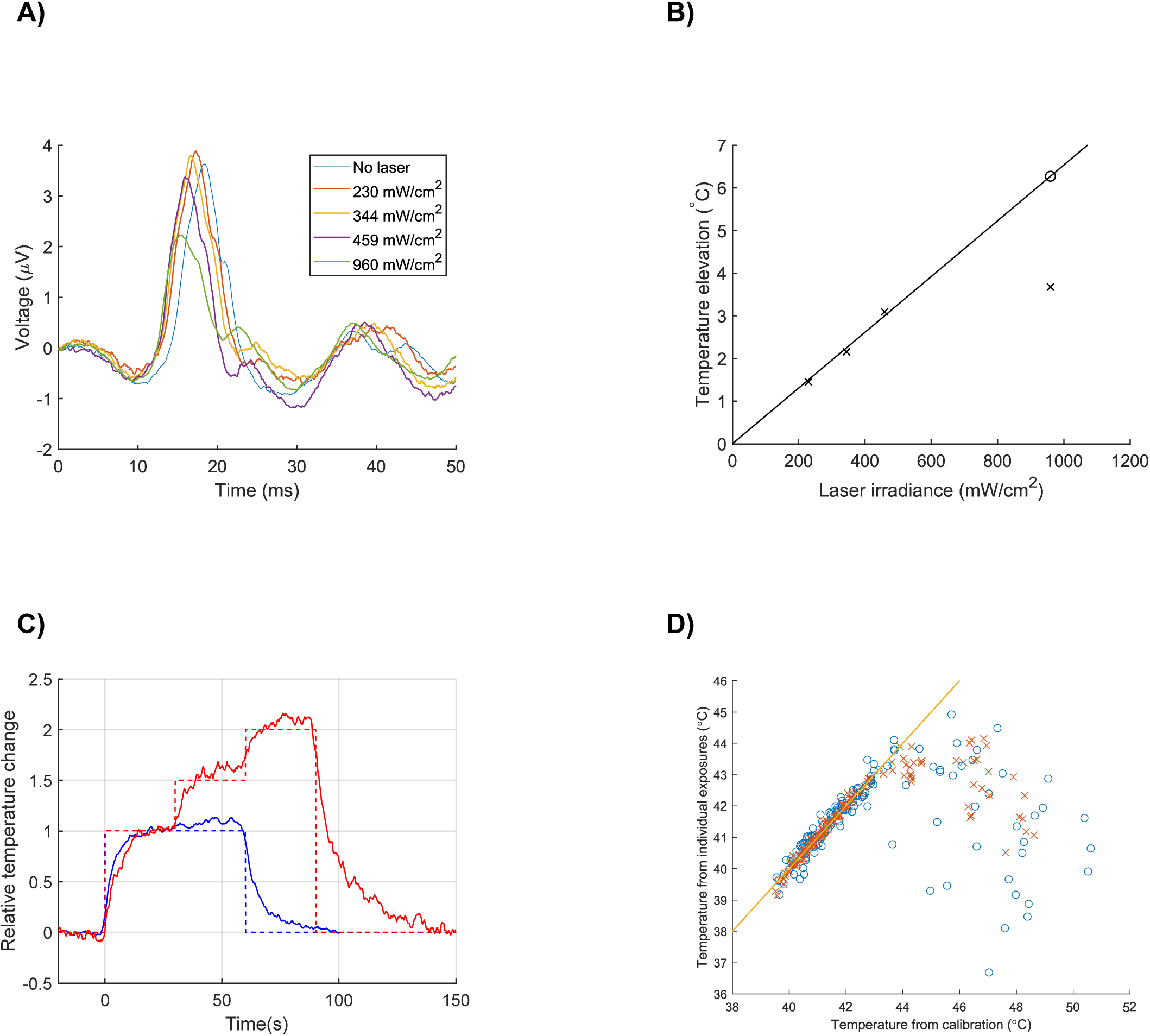
Temperature determination from fERG responses. **A)** Responses acquired without laser application and during a steady-state temperature elevation induced with four different laser powers. **B)** The ERG-based retinal temperature estimate computed from responses of panel A (black crosses) and the target laser power needed to increase the retinal temperature by 6.2 °C (blue circle). In the calibration protocol, the target temperature is extrapolated from the zero intersecting linear fit to the data points gained from three low power laser exposures. Estimate can be converted to absolute temperature by adding the pig body temperature to the determined value. **C)** Normalized retinal temperature elevation averaged across all laser power calibration protocols. The solid blue trace shows the time course of the temperature elevation from 60-second laser exposures with constant power. The solid red trace shows the temperature elevation from three consecutive laser exposures with relative laser powers of 1, 1.5 and 2 (relative laser powers shown with dashed traces). The temperature traces were normalized to produce a temperature elevation of 1 between 15 and 30 seconds after the start of laser application. **D)** linearity of fERG-based retinal temperature determination. Each point corresponds to one laser exposure (crosses for 5 mm spot, circles for 3.4 mm spot). The y-axis shows the retinal temperature determined directly from the fERG response acquired during laser exposure. The x-axis shows the temperature estimated with the calibration protocol explained above.

The kinetics of the fERG signal was found to accelerate linearly with mild laser exposures, which is well in line with former studies showing a linear relationship between the temperature increase and laser power (Cain, Welch and Michie, 1973; Priebe, Cain and Welch, 1975; Ibarra *et al*., 2004; Heussner *et al*., 2014). However, with the highest laser power, the kinetics of the fERG signal accelerated less than what would be expected from linear behavior. To overcome this nonlinear behavior of our temperature estimate with high temperatures, we started each treatment with a calibration protocol comprising three mild laser exposures producing temperature elevations below 5 °C. A zero-intersecting regression line was fitted between the used laser power and the gained temperature elevation to determine the temperature increase per unit of laser power (slope of the black line in Fig. 4 B). The determined thermal responsivity was then used to estimate the laser power required to reach a higher target temperature in the same retinal area (the black circle in Fig 4 B).

Two different laser spot diameters of 3.4 mm and 5 mm were used in the study for laser treatments. With the 3.4 mm laser spot, we always conducted three separate 60-second laser calibration exposures, while with the 5 mm laser spot, we could shorten the calibration procedure to three consecutive 30-second exposures owing to of higher signal-to-noise ratio of fERG signal with a larger spot size. The average time course of the retinal temperature change during these calibration exposures is illustrated in Fig. 4 C. With a constant power laser exposure, the retinal temperature changes rapidly during the first 15 seconds followed by a very slow upward drift. In further analysis, the fERG responses acquired during laser exposures were processed from the fERG trace recorded during the laser exposure excluding the first 15 seconds for the response to better reflect the peak temperature increase induced by the laser exposure.

Fig. 4 D shows the behavior of the fERG signaling kinetics in a broad temperature range. The temperature estimates followed retinal temperature close to 44 °C, but above 45 °C the ERG kinetics started to decelerate with rising temperatures. The average power required to induce one degree of temperature elevation was 25 ± 7.6 mW (range, 16…51 mW) with 3.4 mm diameter laser spot, and 37 ± 11 mW (range, 21…79 mW) with a 5 mm diameter laser spot, when calculated based on the linear range temperature estimates. The mean thermal responsivity values correspond to 7.4 mW/°C·mm and 7.3 mW/°C·mm of laser spot diameter with 3.4 mm and 5 mm laser spots, respectively. The data supports the conclusion by Mainster and Reichel (Mainster and Reichel, 2000) that laser power should be changed in proportion to the spot diameter rather than to the spot area to achieve equivalent temperature elevations.

### Focal ERG-controlled transpupillary laser heating does not harm the cornea

Thirty-six 60-second laser treatments were conducted with 3.4 mm spot diameter to 10 eyes of 7 pigs and 36 treatments with 5 mm spot to 8 eyes of 4 pigs. Target temperatures ranged from 43.6 to 50.6 °C and from 43.6 to 48.6 °C with 3.4- and 5-mm laser spots, respectively. The laser power required to achieve the desired temperature was determined with the calibration protocol separately for each treatment spot before the treatment exposure. After the treatments, we investigated the laser safety for the anterior eye by visual inspection and fluorescein staining. None of the pigs showed burning marks or other laser-induced adverse events on the cornea. However, five eyes showed corneal abrasion most likely due to long-lasting and repetitive use of contact fundus lens and ERG electrode. No irritation or alteration in general habitus was detected in any animal during the recovery time of 24.5 ± 1.8 hours (mean ± std) before euthanasia.

### Focal ERG-based retinal temperature determination predicts HSP70 production and lesion generation with high accuracy

Generation of lesions was evaluated with fundus imaging, fluorescein angiography, and color photography from opened pig eyecup (Fig. 5 A-C). A treatment was judged to cause a lesion if retinal whitening was visible in any of the imaging modalities in the treated area. HSP70 production was investigated with fluorescence microscopy by immunostaining of RPE flat mounts prepared from the posterior eyecups of pigs treated with 3.4 mm spot size. Fig. 5 D-F show exemplary immunostainings for treatment to temperatures 45.7, 47.5, and 50.4 °C, respectively (Spots 5, 2, and 4 in Fig. 5 A, respectively. Elevation of HSP70 expression was easily discernible in all treatments and the fluorescent intensity increased with temperature. Moreover, the laser impact on the morphology of RPE cells was examined by using fluorescent-labeled phalloidin to visualize polymerized actin filaments. Treatments to 45.7 and 47.5 °C showed no visible damage or alterations in the morphology or RPE cells, while treatment to 50.4 °C caused a ring-shaped HSP70 expression with particularly significant expression at the edge of the heated area as well as obvious signs of damage in actin staining. The observed ring-shaped is likely associated with low RPE cell survival and increasing apoptotic cell death (Fig. 5 F) (Amirkavei *et al*., 2020). Hence, increasing the temperature above the damage threshold is assumed to increase the damaged area instead of improving therapeutic response via HSP70 overexpression.

**Figure 5.**
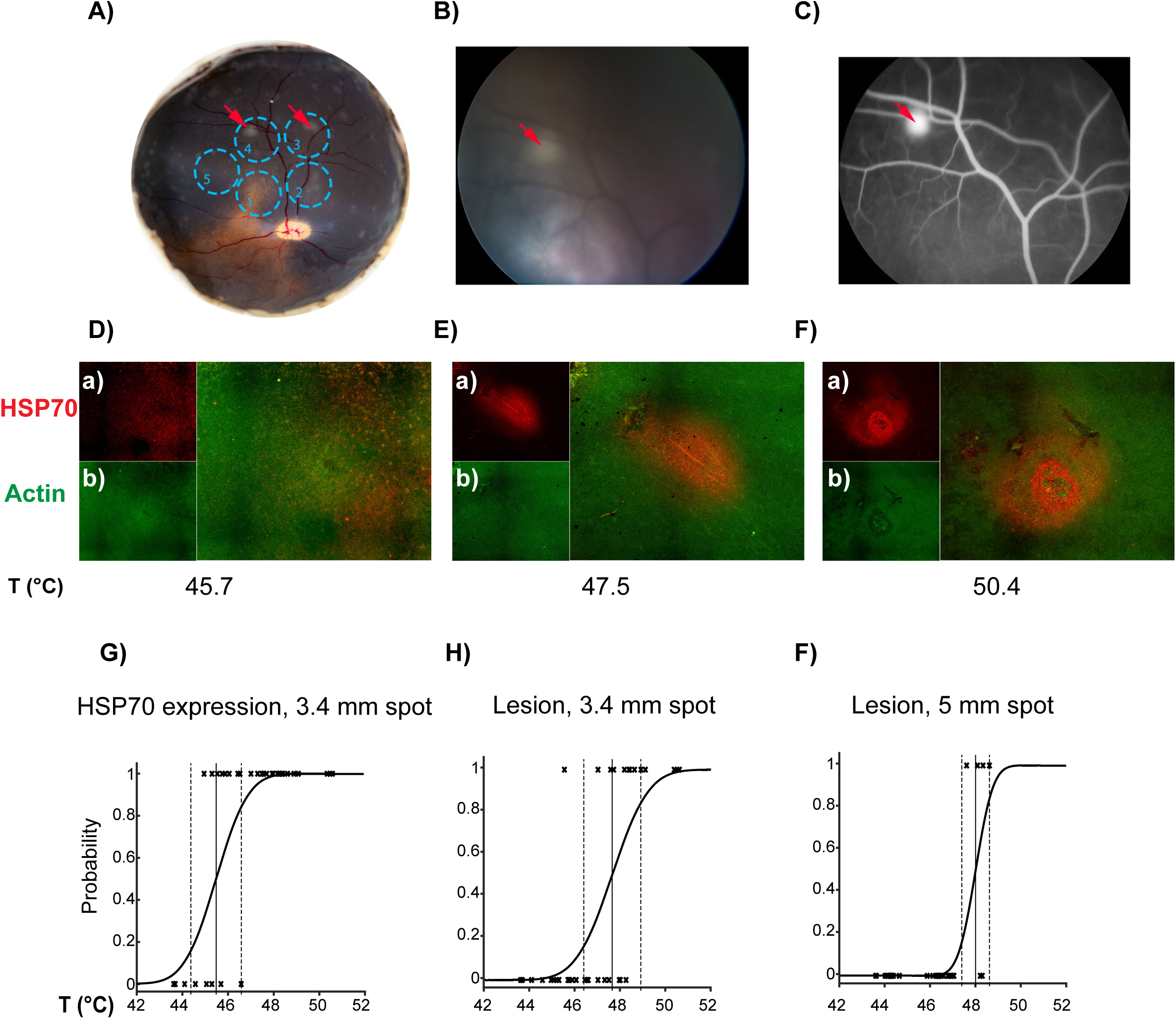
Representative examples for lesion generation and HSP70 expression after SLT across different temperatures. **A)** Image of an open eye cup taken immediately after the isolation of the eye. Blue dashed circles show the placement of 5 consecutive 3.4 mm treatment spots on the fundus. The reached temperatures were 47.3 °C for #1, 47.5 °C for # 2, 48.9 °C for #3, 50.4 °C for #4, and 45.7 °C for #5. **B)** Fundus camera image. **C)** Fluorescein angiography image. The red arrows show the detected lesion from the images. The lesion caused by treatment #3 was visible in the eyecup image but not in fundus or fluorescein angiography images. **D-F)** HSP70 expression in RPE flat-mount across different temperatures. Panels (a) represent HSP70 staining (red); panels (b) actin staining by phalloidin (green). **G-I)** Probability analysis to estimate HSP70 expression threshold for 3.4 mm laser spot and ED50 lesion threshold for 3.4- and 5-mm laser spot across different temperatures. Solid vertical line presents the ED50 values, and the dashed lines presents the 95% confidence intervals for the determined ED50 values.

Each treatment was given a binary classification for triggering HSP70 production and for causing a lesion. A probit model was fitted between the binary classifications and the target temperatures to determine the threshold temperatures for the examined events (Fig. 5 G-I). The determined ED50 temperature for induction of HSP70 production, clearly visible in immunostaining, was 45.5 °C ± (CI 95 %: 44.8 – 46.2 °C) for 3.4 mm laser spot, while the ED50 temperatures for lesion generation were 47.7 °C (CI 95 %: 47.0 – 48.4 °C) and 48.0 °C (CI 95 %: 47.5 – 48.6 °C) with 3.4- and 5-mm spots, respectively.

The probit model was also used to estimate the total accuracy of the thermal dosimetry method to predict lesion generation. This analysis produced standard error values of 1.3 °C (CI 95 %: 0.84 – 3.4 °C) and 0.60 °C (CI 95 %: 0.32 – 1.7 °C) for 3.4- and 5-mm laser spot data, respectively. The respective standard errors relative to caused temperature elevations were 13 % (CI 95 %: 8.8 – 34 %) and 6.7 % (CI 95 %: 3.4 – 18 %).

### Drop in the focal ERG signal amplitude precedes lesion generation

Fig. 6 shows the amplitude change of the fERG responses for 5 mm spots collected during laser exposure at peak temperature and the recovery of the amplitude after the retina returns to normal body temperature. The amplitude did not change drastically at 44 °C but dropped precipitously when the temperature was increased further. After termination of non-damaging laser exposures, the signal amplitude typically recovered near pre-treatment levels. However, in treatments producing a lesion, the fERG signal amplitude first reduced to 32 ± 11 % (mean ± std) but recovered only to 67 ± 8.8 % (mean ± std) after the treated area had returned to body temperature.

**Figure 6.**
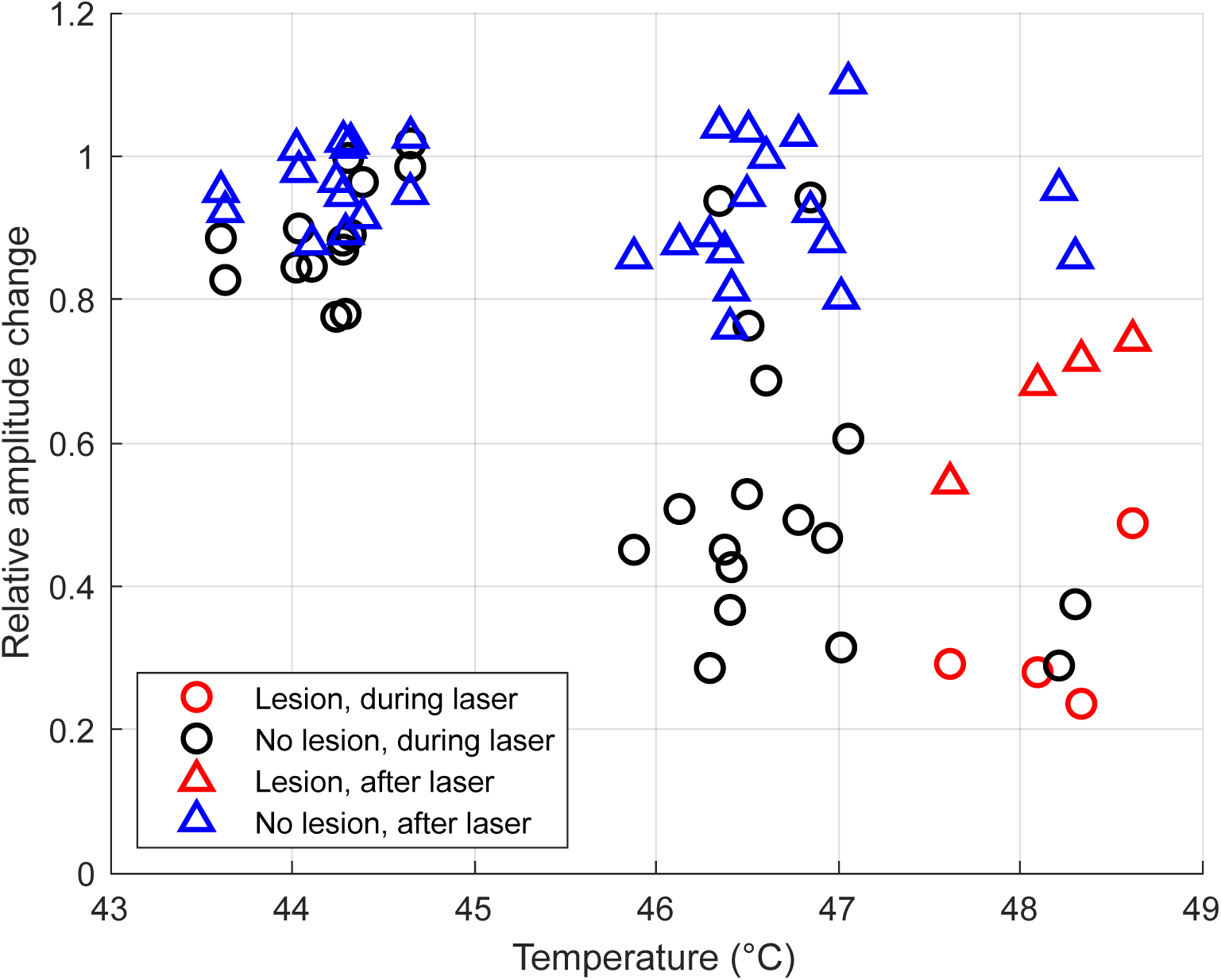
Effects of retinal hyperthermia on fERG response amplitude. Relative changes in fERG amplitudes during (black circles) and after the laser exposure (blue triangles) compared to fERG amplitudes recorded before the laser exposure. Treatments that caused lesions are highlighted with red. fERG amplitudes during the treatment are acquired 30 to 60 seconds after the initiation of laser exposure while the responses after the laser exposure are collected from 30 to 60 seconds succeeding the laser turn-off. The x-axis shows the steady-state temperature of each laser exposure determined with the ERG-based laser power calibration protocol.

### Subthreshold laser exposure induces changes in gene expression in the RPE/choroid

The laser treatments with 5 mm spot size were chosen for further molecular analysis. The treatment spots were divided into three temperature groups, with average target temperatures of 44.2 °C (std = 0.3 °C, n = 10), 46.5 °C (std = 0.3 °C, n = 12) and 48.2 °C (std = 0.3 °C, n = 5), based on the probability distributions for causing visible damage (47.7 – 48.0 °C) and for triggering HSP70 production visible in immunostainings (45.5 °C) (Fig. 5 G-I). The target temperature for the lowest group was selected to have a low probability of triggering HSP70 production observed in immunostaining and a negligible probability of causing visible damage while the middle group had a high probability of triggering HSP70 production and a low probability of damage. The highest group was used as a positive control for damage.

The changes in gene expression induced by subthreshold laser exposure are shown by the Volcano plots in Fig. 7. The complete set of genes studied is given in Supplementary Table 1. The general impression of the figure is that significant upregulation was more common than downregulation and that upregulation was more substantial at 44.2 °C (Fig. 7 A) compared to 46.5 °C (Fig. 7 B). Further, a large part of the upregulated genes consists either of heat shock genes or of autophagy genes.

**Figure 7.**
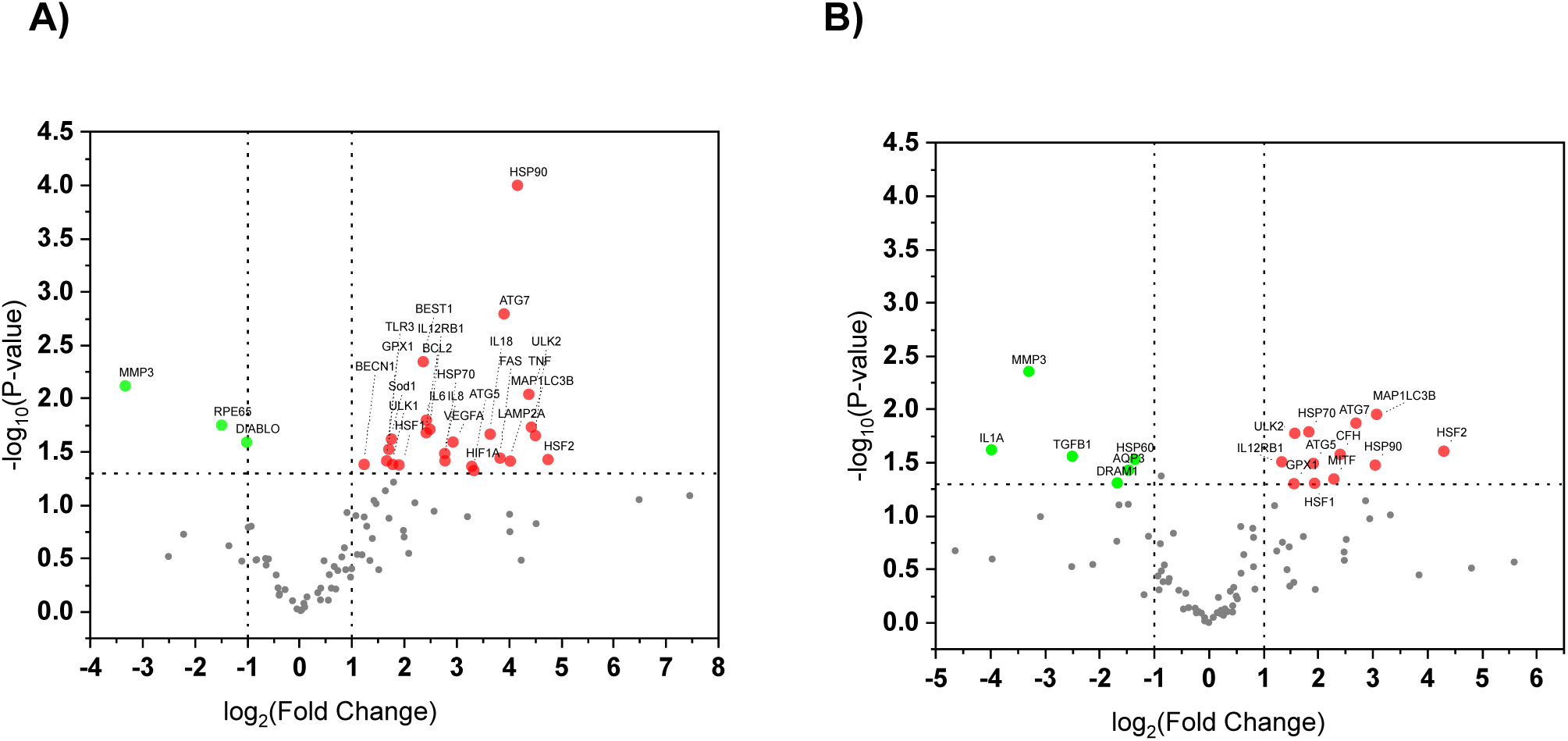
Volcano plots showing the differential gene expression profiles of 91 genes in RPE/choroid cells. Gene expression of cells 24 h after SLT was shown at different temperatures of **A)** 44.2 °C and **B)** 46.5 °C. The −log10 (p-value) was plotted against log2 (fold change (treatment/non-treated control)). Each dot represents the average value of one gene. Genes with significantly differential expression (p-value < 0.05) are highlighted in red (upregulated) and green (downregulated). N= 10 for non-treated control, N=10 for 44.2 °C, N=12 for 46.5 °C.

### Subthreshold laser exposure induces heat shock response in the RPE/choroid

We first examined the changes in heat shock factor and heat shock protein gene expressions in response to SLT in the 5 mm laser-treated spots of the RPE/choroid. Fig. 8 A shows that the gene expression of *HSF1, HSF2, HSP90* and *HSP70* were significantly upregulated for both 44.2 and 46.5 °C. The SLT-induced increase in expression was higher at 44.2 °C than at 46.5 °C for all these genes but HSF1. However, the gene encoding the mitochondrial chaperone *HSP60* was significantly downregulated by SLT at 46.5 °C. The SLT-induced changes in protein levels were evaluated by immunoblot analysis for both RPE/choroid and the neural retina. The HSF1 protein level did not show any changes for the examined temperatures (Fig. 8 B). This could indicate that during the recovery time until euthanasia, both HSF1 expression and post-translation modifications were returned to their non-treated control (NTC) levels, in agreement with the HSF1 behavior observed in cultured ARPE-19 cells (Amirkavei *et al*., 2022). SLT at 44.2 °C did not change the HSF2 protein level, but it was significantly increased for the 46.5 °C (*p < 0.05 vs NTC, &&p < 0.01 vs 44.2 °C; Fig. 8 B). SLT at 44.2 °C approximately doubled the level of HSP90 and HSP70 proteins for both temperatures in RPE/choroid cells (* p < 0.05 vs NTC, ***p < 0.001 vs NTC, Fig. 8 C) and further doubled the level with 46.5 °C (*** p < 0.001 vs. NTC, &&& p < 0.001 vs. 44.2 °C, Fig. 8 C). This difference in results is likely due to the higher accuracy of immunoblot analysis compared to the immunostaining method. Our immunoblot analysis demonstrated that SLT did not change the amount of HSP70 in neural retinal cells (Supplementary Fig. 1). However, the HSP90 was significantly increased in neural retinal cells at 46.5 °C (**p < 0.01 vs NTC, &&& p < 0.001 vs 44.2 °C, Supplementary Fig. S1 A).

**Figure 8.**
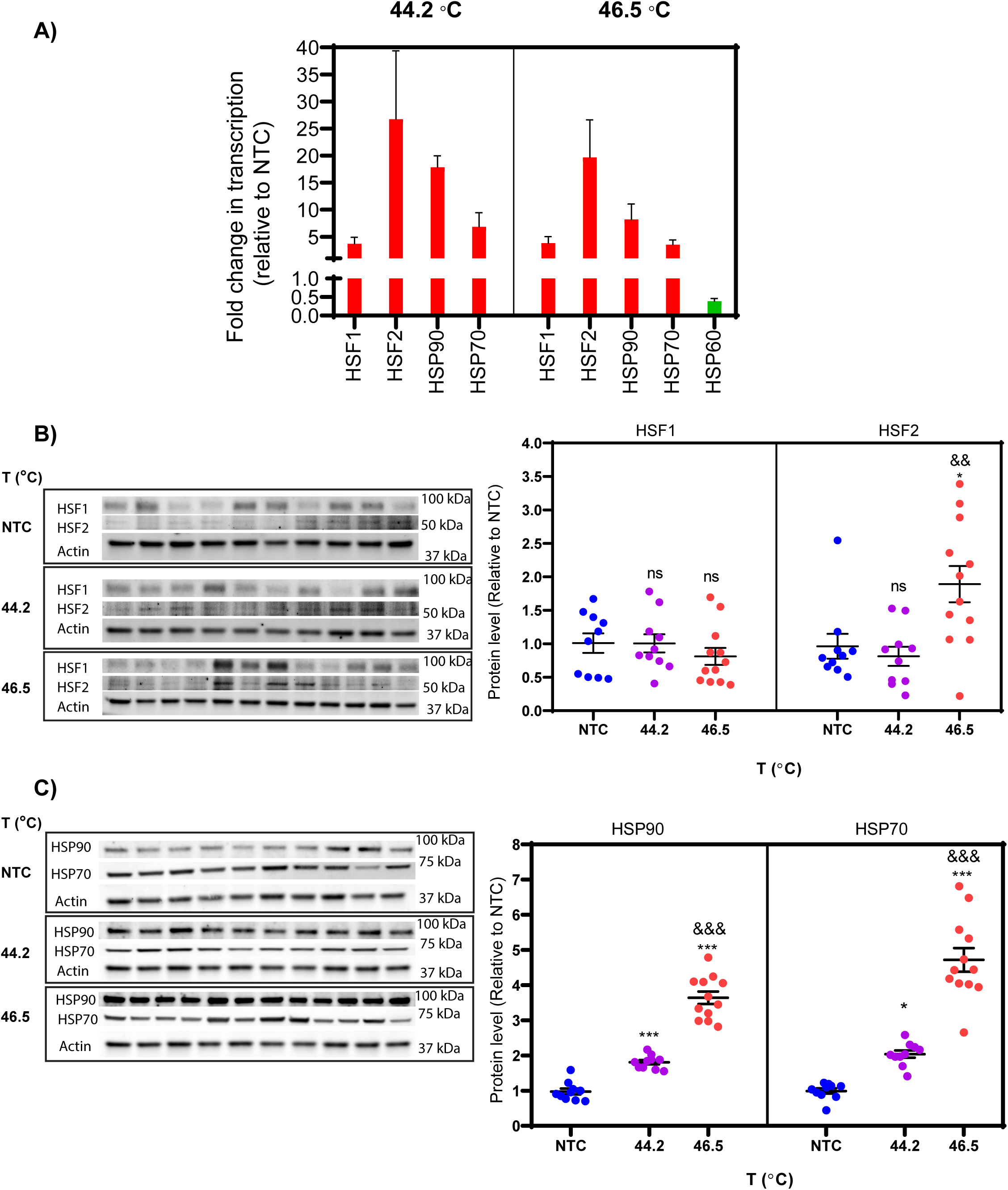
Heat shock induces heat shock response in RPE/choroid cells. **A)** The fold changes in the expression level of heat stress-related transcripts in RPE/choroid at 44.2 °C and 46.5 °C were considered significant with a p-value < 0.05 (data obtained from Fig.7). Cell lysates were analyzed by immunoblot analysis for **B)** HSF1 and HSF2 and **C)** HSP90 and HSP70. Quantitative analysis of protein level was corrected to Actin as the loading control. NTC stands for non-treated control. Data are represented as mean ± SEM (n = 10 for NTC, n=10 for 44.2 °C, n=12 for 46.5 °C). Statistical analysis was used by one-way ANOVA followed by Bonferroni‘s multiple comparisons (* p < 0.05 vs. NTC, *** p < 0.001 vs. NTC, && p < 0.01 vs. 44.2 °C, and &&& p < 0.001 vs. 44.2 °C).

### Heat activates autophagy in the laser-treated RPE/choroid spots

Previously we have demonstrated that autophagy can be activated by a 30 min 42 °C heat shock in cultured ARPE-19 cells (Amirkavei *et al*., 2022). To evaluate whether activation of autophagy can be achieved in RPE/choroid *in vivo* by our temperature-controlled SLT, we first assessed SLT-induced changes in the expression of a selected group of autophagy-associated genes by qPCR. Fig. 9 A presents the changes at 24 h after the SLT for 60 s treatments raising the temperature to 44.2 or 46.5 °C, respectively, in the treated 5 mm spots. SLT increased the expression of many autophagy-associated genes, and the increase was more pronounced at 44.2 than at 46.5 °C. Our data demonstrate that the key genes involved in autophagy induction (*ULK2*), autophagosome formation (*ATG5, ATG7, LC3B*), as well as lysosomal fusion (*LAMP2*), were upregulated at 44.2 °C more than ten-fold, while the expression of the genes *ULK1* (autophagy induction) and *BECN1* (phagophore expansion) were less but still significantly increased (Fig. 9 A). With SLT to 46.5 °C, the increase in expression was smaller than with SLT to 44.2 °C, suggesting that 46.5 °C may exceed the optimal SLT temperature regarding autophagy activation. Consistent with this, the genes DNA damage regulated autophagy modulator 1 (*DRAM1*), which mediates autophagy and cell death, and the transforming growth factor (*TGFB1*), respectively, were significantly downregulated at 46.5 °C (Fig. 9 A).

**Figure 9.**
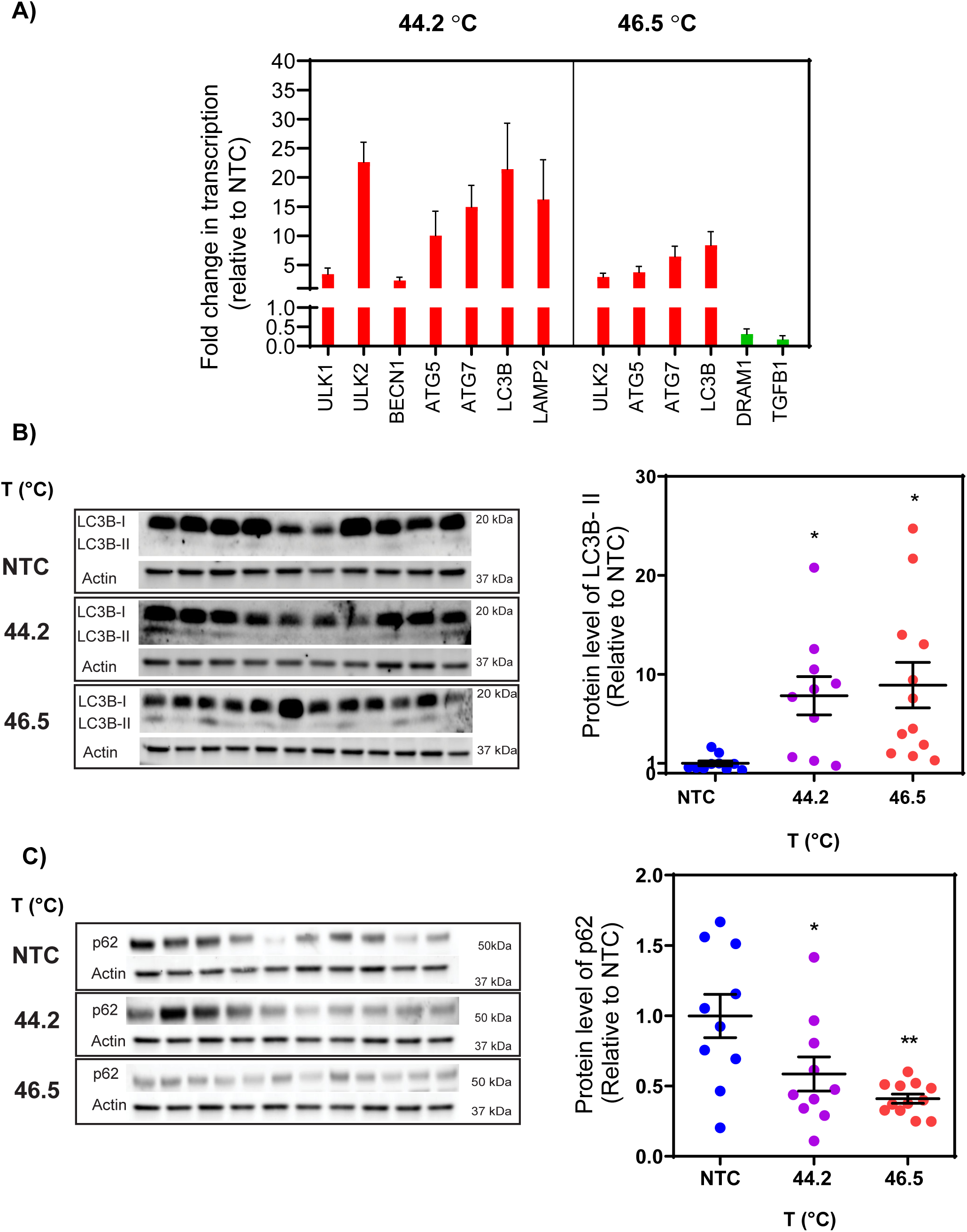
Heat shock regulates induction of autophagy in RPE/choroid cells. **A)** The fold changes in the expression level of autophagy-related transcripts in RPE/choroid were considered significant at 44.2 and 46.5 °C with a p-value < 0.05 (data obtained from Fig.7). Cell lysates were analyzed by immunoblot analysis for **B)** LC3-II and **C)** p62. Quantitative analysis of protein level was corrected to Actin as the loading control. NTC stands for non-treated control. Data are represented as mean ± SEM (n = 10 for NTC, n=10 for 44.2 °C, n=12 for 46.5 °C). Statistical analysis was used by one-way ANOVA followed by Bonferroni‘s multiple comparisons (* p < 0.05 vs. NTC, ** p < 0. 01 vs. NTC).

To investigate whether the elevated autophagy gene expression leads to changes in protein levels, we determined the protein levels of LC3B and p62, two canonical autophagy proteins vital for autophagosome formation in the elongation step. Immunoblot analysis demonstrated a significant increase in the LC3B-II level (* p < 0.05 vs NTC, Fig. 9 B) and a significant decrease in the p62 protein level (* p < 0.05 vs NTC, ** p < 0.01 vs NTC, Fig. 9 C) in RPE/choroid cells at both temperatures of 44.2 and 46.5 °C. These results are consistent with the earlier observations in *C. elegans* and in ARPE-19 cells, where autophagy activation led to elevated LCB-II and lowered p62 levels (Kumsta *et al*., 2017; Amirkavei *et al*., 2022). No changes in the LC3B-II and p62 protein levels were observed in neural retina cells (Supplementary Fig. S1 B & C). Altogether, the data suggest that SLT enhanced autophagy activity in RPE/choroid cells *in vivo*.

### HSP70 hyperexpression and autophagy activation can be triggered by temperature-controlled SLT without causing apoptosis or oxidative stress

To evaluate whether our SLT induces oxidative stress, we first examined the effect of laser treatment to the expression of six oxidative stress-related genes (glutathione peroxidase, *GPX1;* glutathione synthetase *GSS*; heme oxygenase 1, *HMOX1*; nitric oxide synthase 2, *NOS2*; and superoxide dismutases 1 and 2, cytosolic *SOD1* and mitochondrial matrix *SOD2*, respectively). Out of these, only the major antioxidant enzyme genes *GPX1* and *SOD1* showed significant upregulation at 44.2 °C. Further, *GPX1* overexpression alone was significant at 46.5 °C (Fig. 10 A). Next, we measured the total ROS content in the tissue homogenate of RPE/choroid and neural retina separately using the DCFH-DA assay (Fig. 10 B). To provide a positive control group, we analyzed an extra group of samples where a lesion occurred (mean temperature of 48.2 °C). No significant change in ROS content was observed for RPE/choroid and neural retina cells at 44.2 and 46.5 °C compared to NTC, though the slightly higher ROS level observed at 46.5 °C compared to 44.2 °C was significant (& p < 0.05 vs 44.2 °C, Fig. 10 B). The ROS content increased significantly at 48.2 °C compared to NTC, 44.2 °C, and 46.5 °C for both RPE/choroid and neural retina cells (***p < 0.001, and **p < 0.01 vs NTC, Fig. 10 B). These results together indicate a clear activation of oxidative stress response at 48.2 °C but not at 44.2 and 46.5 °C.

**Figure 10.**
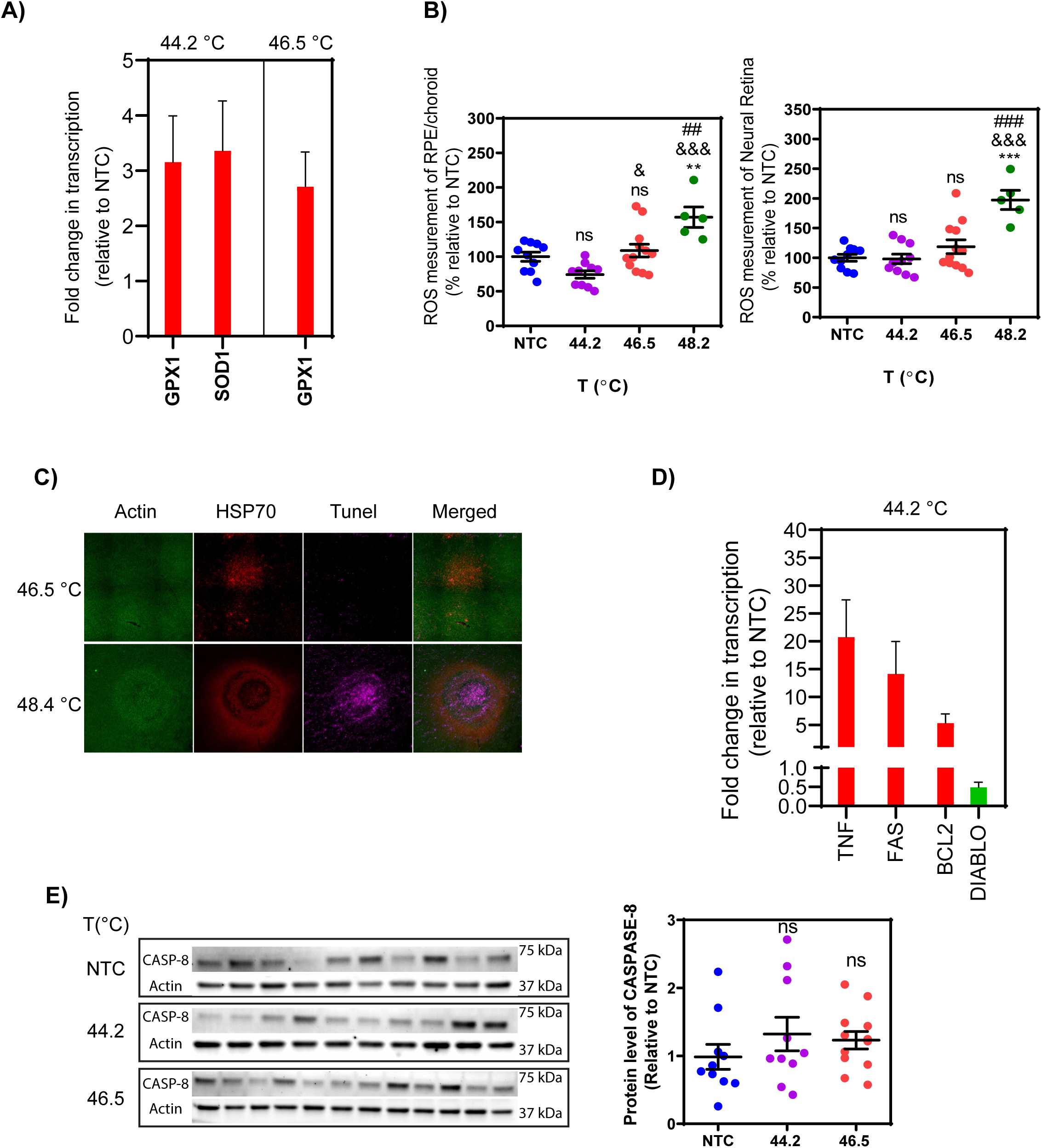
No changes in oxidative stress and apoptosis level after SLT in RPE/choroid and neural retinal cells. **A)** The fold changes in the expression level of oxidative stress-related transcripts in RPE/choroid were considered significant at 44.2 °C with a p-value < 0.05 (data obtained from Fig.7). **B)** Percentage of ROS level for RPE/choroid and neural retinal cells after laser irradiation at 44.2 and 46.5 °C compared with the NTC cells. **C)** Tunel staining of RPE flat mount for laser irradiation at 46.5 and 48.4 °C. Panels (red) represent HSP70 staining, panels (green) display actin staining by phalloidin, and panels (magenta) Tunel staining. **D)** The fold changes in the expression level of apoptosis-related transcripts in RPE/choroid were considered significant at 44.2 with a p-value < 0.05. **E)** Immunoblot analysis of CASP8 for RPE/choroid cells is illustrated with representative images. Quantitative analysis of protein level was corrected to Actin as the loading control and normalized to NTC. NTC stands for non-treated control. Data are represented as mean ± SEM (n = 10 for NTC, n=10 for 44.2 °C, n=12 for 46.5 °C). Statistical analysis was used by one-way ANOVA followed by Bonferroni‘s multiple comparisons (** p < 0.01 vs. NTC, & p < 0. 05 vs. 44.2 °C, &&& p < 0. 001 vs. 44.2 °C, ## p < 0. 01 vs. 46.5 °C, ### p < 0. 001 vs. 46.5 °C).

To evaluate possible RPE cell damages caused by our SLT, we used Tunel assay for RPE flat mounts with 3.4 mm laser spots 24 h after irradiation. The Tunel staining did not detect any marks for apoptosis for the treatments below 48 °C. Therefore, we chose to demonstrate the staining corresponding to the temperature of 46.5 and 48.4 °C as examples for negative and positive staining (Fig. 10 C, panel Tunel). Noticeable apoptotic staining in the middle of laser-treated area at 48.4 °C indicates severe injury in RPE cells. In addition to Tunel assay, we studied the expression of a set of apoptosis-related genes. Heating to 44.2 °C (5 mm spots) caused significant upregulation of tumor necrosis factor *(TNF*, promoter of cell survival and the most potent inducer of apoptosis), *FAS* (death receptor belonging to the *TNF* receptor superfamily), and *BCL2* (Bcl-2 is a mitochondrial outer membrane protein that blocks the mitochondrial apoptotic pathway) in RPE/choroid cells (Fig. 10 D). A significant downregulation of *DIABLO (*a protein that eliminates the inhibitory effect of IAPs (members of the inhibitor of apoptosis protein family)) was observed at 44.2 (Fig. 10 D). Although we did not observe significant changes in *CASP8*, coding the caspase-8 (CASP8) protein which has a pivotal role in the extrinsic apoptotic signaling pathway and is involved in the programmed cell death induced by FAS. Consistent with our Tunel staining findings, the immunoblot analysis did not demonstrate any significant difference in the expression of CASP8 for both temperatures of 44.2 and 46.5 °C compared to NTC in RPE/choroid cells (Fig. 10 E). CASP8 level for neural retinal cells was not different for both temperatures compared to NTC, but CASP8 increased significantly at 46.5 °C compared to 44.2 °C (& p < 0.05 vs 44.2 °C, Supplementary Fig. S2 A). This finding could be related to tissue-specific variation in thermotolerance in RPE/choroid and neural retinal cell and highlights the importance of knowing temperature during SLT.

### SLT modulates VEGF and RPE survival factors

Strong laser treatment may act as a local insult, and one possible source of tissue damages might be vascular destabilization, which may progress to vasculogenesis, angiogenesis, and ultimately to neovascularization. Data for gene expression demonstrated a significant upregulation of *HIF1A* and *VEGF A* genes at 44.2 °C in the RPE/choroid cells (Fig. 7 A). Subsequently immunoblot analysis was conducted to evaluate the secretion of VEGF165, the major isoform coded by *VEGF A* (Fig. 11). The protein level analysis of VEGF165 did not show any change on its expression at 44.2 °C in RPE/choroid cells, while a significant increase in expression of VEGF165 was found at 46.5 °C (* p < 0.05 vs NTC, Fig. 10 B). The analysis of VEGF165 protein in neural retina demonstrated a significant decrease at 44.2 °C, with no discernible secretion at 46.5 °C (Supplementary Fig. S2 B).

**Figure 11.**
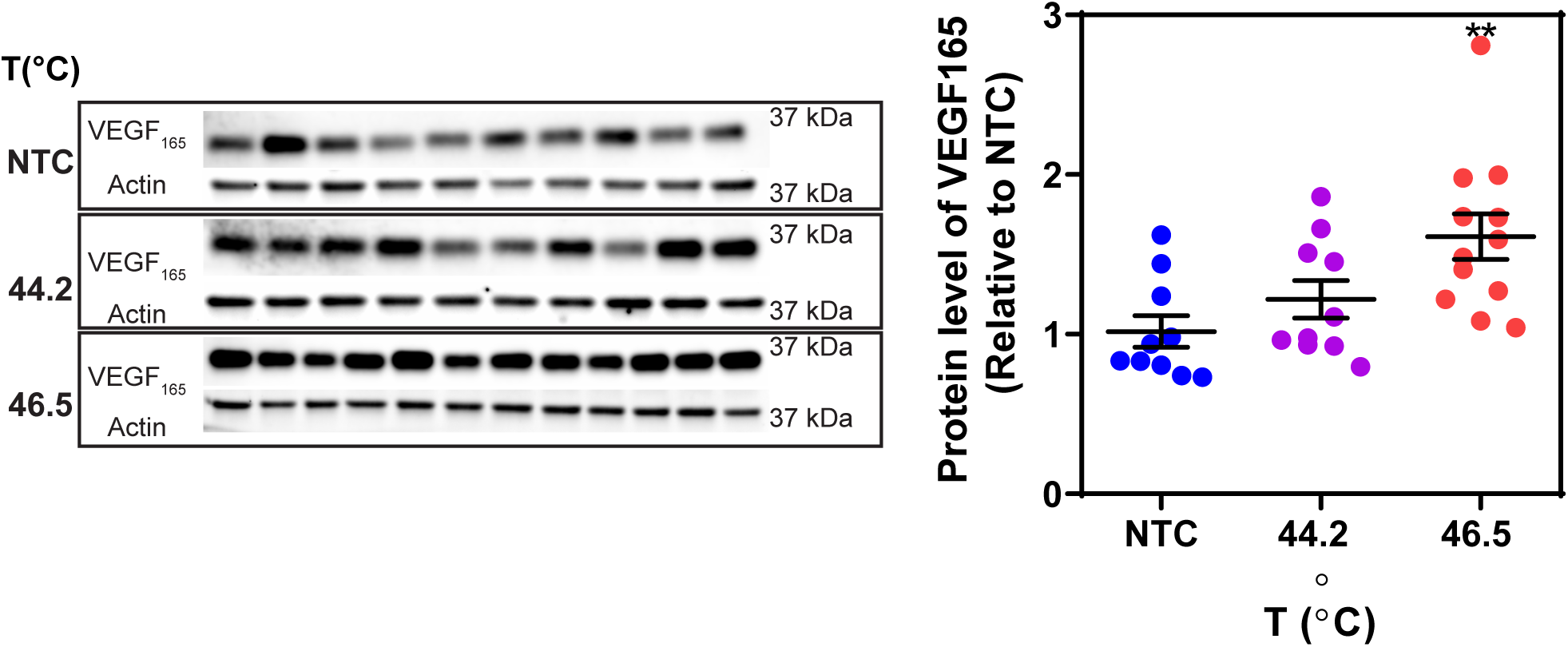
Induced heat shock doesn’t cause VEGEF-mediated angiogenesis in RPE/choroid cells. Immunoblot analysis for VEGF-165 for RPE/choroid cells is illustrated with representative images. Quantitative analysis of protein level was corrected to Actin as the loading control and normalized to NTC. Data are represented as mean ± SEM (n = 10 for NTC, n=10 for 44.2 °C, n=12 for 46.5 °C). Statistical analysis was achieved using one-way ANOVA, followed by Sidak test (** p < 0.01 vs. NTC).

## Discussion

Protein homeostasis (proteostasis) is essential for the functioning of proteins in all living cells. Proteostasis is maintained by a complex proteostasis network controlling protein synthesis, folding, trafficking, aggregation, and degradation. Many diseases appear to be caused by improper proteostasis, including the gain-of-toxic-function diseases like Alzheimer’s, Parkinson’s, and Huntington’s disease, as well as AMD (Chung, Dawson and Dawson, 2001; Algvere and Seregard, 2002). With aging, all the major protein quality control mechanisms, i.e. protein folding and misfolded protein refolding by molecular chaperones, protein degradation by the ubiquitin-proteasome system, and degradation of aggregates and dysfunctional cell organelles by autophagy, are known to lose their effectiveness (Viiri *et al*., 2013; Mitter *et al*., 2014). Malfunctioning of the protein quality control in the postmitotic RPE cells is believed to play a pivotal role in the generation and progression of AMD, manifested as the accumulation of waste material and drusens in the RPE cells and the Bruch’s membrane (Kaarniranta *et al*., 2011). Continuous high oxidative load and decreased capacity of aged cells to cope with cellular stresses cause dysfunction of RPE cells, breakdown of photoreceptors, and ultimately a loss of central vision in AMD patients (Bhutto and Lutty, 2012). In this study, we demonstrate in pigs *in vivo* a novel SLT modality implementing a fERG-based thermal dosimetry that allows controlled activation of two of the key protein quality control mechanisms, the molecular chaperone system (heat shock proteins) and autophagy. With our method, a 5 mm spot in the pig fundus, corresponding to the 5.5 mm diameter human macula, was heated with a near-IR laser while simultaneously monitoring the temperature of the heated spot with an error less than 0.6 °C. The method allowed us to investigate the temperature window of SLT-induced effects and to find a temperature window, which induced heat shock protein production and activated autophagy in RPE/choroid without signs of oxidative stress, apoptosis, or morphological changes in the RPE/choroid or neural retina. Our data suggests that the optimal heat treatment temperature for HSP and autophagy activation may be close to 44 °C in pigs for 60-second laser exposure.

Our temperature determination expects logarithmic change in the kinetics of photopic fERG signal from the heated area to be linearly dependent on the retinal temperature (Kaikkonen *et al*., 2021). Our data confirmed this expectation to hold up to around 44 °C, after which the response kinetics accelerated less than predicted by a linear model (see Fig. 4 D). The reason for the breakage of the linearity was not studied here, but a typical feature for biological processes is to accelerate towards higher temperatures and decelerate after surpassing the functional temperature range of the process (Cossins and Bowler, 1987). As long as the retinal temperature remains in the linear zone, temperature determination can be conducted solely by the real-time ERG-based temperature determination, while at temperatures beyond the linear zone, temperature should be estimated by the calibration protocol demonstrated in the study. Additionally, the fERG signal amplitudes reduced significantly after surpassing 44 °C (Fig. 6). The nonlinear behavior of fERG kinetics and amplitude drop could potentially be used as independent safety features implying an overheating in treatments close to 44 °C target temperature.

With the introduced calibration protocol and 60-second treatments using 5 mm laser spot size, the RMS temperature determination error for the ERG-based thermal dosimetry was 0.6 °C for detecting the temperature generating visible lesions. The presented error includes variation from three main sources: error in the body temperature determination, error in the estimate for retinal temperature elevation, as well as a natural deviation in the damage threshold between individuals. The data cannot separate these sources of error but shows the upper bound for the accuracy of the temperature estimate.

In testing the potentially beneficial effects of our temperature-controlled SLT modality, we focused on two key goals, increasing the HSP70 expression and activation of macroautophagy. Since it was not feasible to make time series analysis on gene expression and protein level changes with pigs *in vivo*, we had previously conducted a more complete set of experiments on cultured human ARPE-19 cells (Amirkavei *et al*., 2022) that allowed comparison with the present data. While in the ARPE-19 cell study we investigated the time behavior (time points from 3 h to 24 h recovery) of gene and protein expression to an identical heat shock (30 min at 42 °C), here we focused on the effects of varying heating temperature at a single time point (24 h recovery). Our data demonstrates that very similar increases in HSF and HSP as well as macroautophagy gene expression can be achieved with 60 s heating sessions in pigs *in vivo* compared to those observed in ARPE-19 cells with 30 times longer heat shocks. Further, the local heating and the fERG-based temperature determination from the heated area of the pig RPE appeared to work reliably, enabling accurately controlled temperature elevations in the heating spot. In ARPE-19 cells, the signs of post-translational modification seemed to be completely over at 12 h post-heat shock (our next time point after the 3 h after recovery time point) (Amirkavei *et al*., 2022), while in pigs the heat-induced HSF1 transcription level was still up (ca. four-fold) at 24 h recovery compared to control. This was, however, not reflected in the total HSF1 protein level. Unfortunately, we did not study post-translation modification of HSF1, because based on the earlier ARPE-19 cell data we expected all the stages of HSF1 activation mechanism be over at 24 h recovery. However, this result may suggest that the HSF1 activation may last longer *in vivo* compared to cultured RPE cells.

In investigating macroautophagy, we focused on two key autophagic cargo recognition proteins involved in selective autophagy, p62 and LC3B-II. A significant increase in the expression of the LC3B-II gene and protein levels combined with a remarkable decrease in the p62 protein level were observed at both temperatures (Fig. 9). This behavior was consistent with our previous results from ARPE-19 cells, where the increase in LC3B-II level and simultaneous decrease in p62 level were shown to be caused by heat shock-activated autophagy. Thus, even without being able to study the effect of heating on autophagic flux in pigs *in vivo*, we conclude that our results on LC3B-II and p62 together with the strongly increased expression of the other key studied autophagy genes (Fig. 9 A) most likely reflect heat-induced activation of selective macroautophagy. Since the same autophagy proteins are involved also in non-selective macroautophagy, it is likely that the non-selective macroautophagy is activated as well. Further, the increased expression of the LAMP2 gene, LAMP2 being a key receptor protein in chaperone-mediated autophagy, suggests that CMA may also be activated by heat.

It is well known that both heat shock proteins and autophagy proteins are under diurnal control and that the gene activation and protein levels vary during the day and night (Sandström *et al*., 2009; Wang *et al*., 2020). Our choice of controls removed the possibility that the observed heat-induced expressions could be due to diurnal variation. In this study we did not either investigate the effect of diurnal control on the level of gene expression and protein synthesis, however, it may be worth noticing that it might be, at least in principle, to optimize the effectiveness of heat-induced HSP and autophagy protein production according to the time of day when the heat treatment is given.

Oxidative stress is tightly linked to the pathophysiology of AMD (Dong *et al*., 2009; Mitter *et al*., 2014). Increased production of antioxidant defense system components has been considered as a promising approach for treating retinal degenerations in which oxidative damage plays a significant role (Iwami 2014). Studies of mice deficient in SOD1 have shown a high basal level of oxidative damage and increased vulnerability to retinal degeneration when exposed to oxidants (Lu *et al*., 2009). Further, SOD1^-/-^ mice develop many characteristics of patients with AMD (Dong *et al*., 2009). In this study, we observed significant heat-induced upregulation of the antioxidant genes coding the enzymes *SOD1* and *GPX1* in RPE/choroid cells (Fig. 10 A). SODs convert superoxide to oxygen and hydrogen peroxide, whereas GPX and catalase degrade hydrogen peroxide into oxygen and water. We expect that the remarkable upregulation of *SOD1* and *GPX1* observed after SLT to 44.2 °C in RPE/choroid cells would improve protection against oxidative stress and help in ROS detoxification. The observation that ROS content in RPE/choroid cells did not increase or rather showed signs of decrease at 44.2 °C suggests that our SLT activates the antioxidant defense system (Fig. 10 B).

Excessive heating may lead to apoptosis. In our study, SLT significantly upregulated (*TNF, FAS, BCl2*) and downregulated (*DIABLO*) apoptosis-related genes at 44.2 °C (Fig. 10 D). These changes suggest that the observed upregulations at 44.2 °C might rather be protective than apoptosis-inducing. TNF-α is known to induce apoptosis in many cell types. However, it also activates autophagy, thereby balancing apoptosis and autophagy (Zheng *et al*., 2017) BCL-2 is an apoptosis regulator, being able to act as either anti- and pro-apoptotic factor. In human RPE cells, the overexpression of *BCl2* has been shown to increase survival cells exposed to oxidative stress induced by H_2_O_2_ (Godley et al., 2002 Exp. Eye Res) suggesting a protective role for BCL-2 in our SLT study. The observation that the gene of the death receptor FAS was upregulated only at 44.2 °C but not at 46.5 °C is somewhat surprising and may suggest that also FAS might have some other role than purely apoptosis-activating. This hypothesis is supported by our TUNEL assay (Fig. 10 C) and CASP8 (Fig. 10 E) results that did not detect cell death for either 44.2 °C or 46.5 °C. This is in agreement with previous *in vitro* studies on cultured RPE cells, where apoptotic cell death has been found to occur only at temperatures above 46 °C and necrosis above 50 °C (Kern *et al*., 2018).

Previous SLT studies have raised a concern that SLTs might increase the risk of neovascularization for eyes with large drusens (Friberg et al., 2006). Hence, we investigated the treatment-induced changes in the production of VEGF, which regulates the proliferation and density of endothelial cells in the choriocapillaris, extracellular space, and around the Bruch’s membrane. We found an increased production of VEGF165 at 46.5 °C in RPE/choroid cells (Fig. 11 B), which might increase a risk for the formation of unusual blood vessels, as observed in AMD. However, heating to 44.2 °C did not affect VEGF production at RPE/choroid and even decreased it in the neural retina, which again highlights the need for precisely controlled temperature for SLTs.

In summary, we showed that heating the retina to roughly 44 °C with a 60-second laser exposure activated promising therapeutical targets (HSPs, autophagy, antioxidants) in RPE/choroid, with a clear margin to damaging temperatures. According to this study, 44 °C can be safely reached by implementing the introduced fERG-based thermal dosimetry. The increased expression of HSP70, recognized as the sole refolding system for misfolded proteins, promotes cellular survival and activation of the antioxidant defense systems, while the activation of the autophagy may increase waste clearance in RPE cells. Activating these natural protective cascades can help to rejuvenate aged and diseases cells and potentially help in managing retinal diseases such as AMD and macular edemas.

---

**Figure S1.**
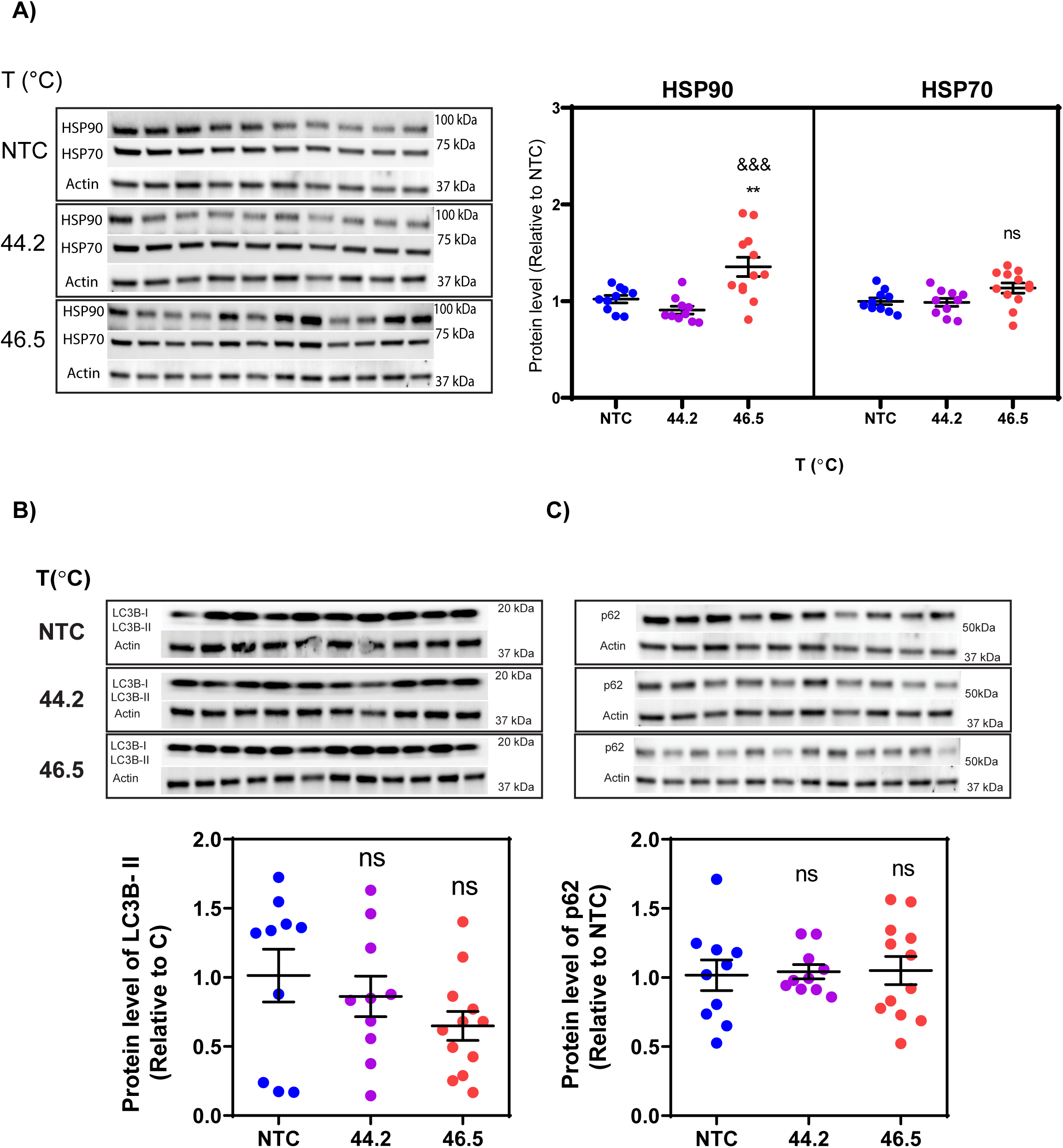
Heat shock induces heat shock response in the neural retina cells and has no effect on their autophagy activation. **A)** Cell lysates were analyzed by immunoblot analysis for HSP90 and HSP70, **B)** for LC3-II and p62. Quantitative analysis of protein level was corrected to Actin as the loading control. NTC stands for non-treated control. Data are represented as mean ± SEM (n = 10 for NTC, n=10 for 44.2 °C, n=12 for 46.5 °C). Statistical analysis was used by one-way ANOVA followed by Bonferroni‘s multiple comparisons (** p < 0. 01 vs. NTC, &&& p < 0.001 vs. 44.2 °C).

**Figure S2.**
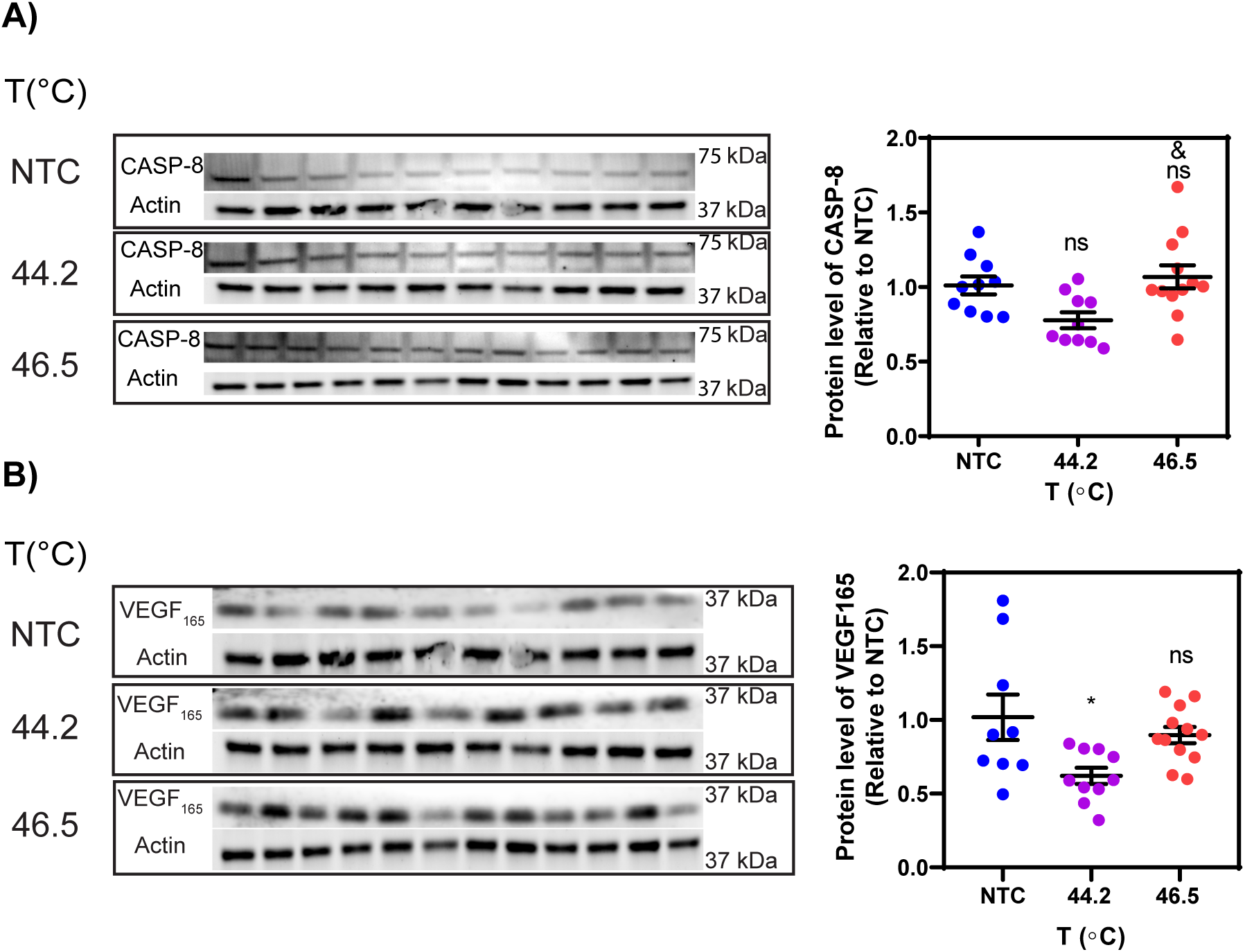
Induced heat shock in the neural retina cells has no effect on the activation of apoptosis while heating the retina to 44.2 °C seems to decrease the VEGF level. **A)** Immunoblot analysis of CASP8 for neural retina cells is illustrated with representative images. **B**) Immunoblot analysis for VEGF-165 for neural retinal cells is illustrated with representative images. Quantitative analysis of protein level was corrected to Actin as the loading control and normalized to NTC. NTC stands for non-treated control. Data are represented as mean ± SEM (n = 10 for NTC, n=10 for 44.2 °C, n=12 for 46.5 °C). Statistical analysis was used by one-way ANOVA followed by Bonferroni‘s multiple comparisons (* p < 0.05 vs. NTC, & p < 0. 05 vs. 44.2 °C).

**Table S1.** Complete list of analyzed genes.

| # | Symbol | Description | GeneBank | Fold change<br>44.2 °C | p-value | Fold change<br>46.5 °C | p-value |
| --- | --- | --- | --- | --- | --- | --- | --- |
| Heat stress |  |  |  |  |  |  |  |
| 1 | CRYAB | Crystallin, alpha B | XM_003357294 | 0,64 | 0,3161 | 0,57 | 0,2872 |
| 2 | HSF1 | Heat shock transcription factor 1 | NM_001243819 | 3,71 | 0,0419 | 3,81 | 0,0491 |
| 3 | HSF2 | Heat shock transcription factor 2 | XM_005659227 | 26,73 | 0,0372 | 19,66 | 0,0246 |
| 4 | HSP90 | Hsp90 co-chaperon cdc37-like 1-like | XM_003121918 | 17,84 | 0,0001 | 8,22 | 0,0332 |
| 5 | HSP70 | Heat shock protein 70 | NM_001123127 | 6,83 | 0,0383 | 3,55 | 0,0162 |
| 6 | HSP27 | Heat shock 27 kDa protein 1 | NM_001007528 | 1,38 | 0,3322 | 1,31 | 0,5076 |
| 7 | HSP60 | Heat shock 60 kDa protein 1 (chaperonin) | NM_001254716 | 0,91 | 0,7884 | 0,39 | 0,0294 |
| 8 | HSP 40 | DnaJ (Hsp40) homolog, subfamily B, member 12 | XM_001924319 | 1,46 | 0,7775 | 0,06 | 0,2534 |
| Autophagy |  |  |  |  |  |  |  |
| 9 | ATG5 | ATG5 autophagy related 5 homolog (S.cerevisiae) | NM_001037152 | 10,02 | 0,0472 | 3,76 | 0,032 |
| 10 | ATG7 | ATG7 autophagy related 7 homolog (S.cerevisiae) | NM_001190285 | 14,93 | 0,0016 | 6,45 | 0,0134 |
| 11 | ATG3 | Autophagy related 3 | XM_003132682 | 18,79 | 0,3286 | 1,23 | 0,7835 |
| 12 | BNIP3L | BCL2/adenovirus E1B 19kDa interacting protein 3-like | XM_001927592 | 174,78 | 0,0813 | 0,72 | 0,7457 |
| 13 | BECN1 | Beclin 1, autophagy related | NM_001044530 | 2,34 | 0,0414 | 0,88 | 0,7874 |
| 14 | DRAM1 | DNA-damage regulated autophagy modulator 1 | NM_001190286 | 3,95 | 0,1724 | 0,31 | 0,0489 |
| 15 | DRAM2 | DNA-damage regulated autophagy modulator 2 | NM_001243696 | 4,59 | 0,095 | 2,77 | 0,1945 |
| 16 | LAMP2A | Lysosomal-associated membrane protein 2 | NM_001244255 | 16,23 | 0,0385 | 0,53 | 0,4927 |
| 17 | LAMP-1 | Lysosomal-associated membrane glycoprotein 1 | NM_001011507 | 1,32 | 0,6 | 1,06 | 0,8932 |
| 18 | MAP1LC3A | Microtubule-associated protein 1 light chain 3 alpha | NM_001170827 | 0,66 | 0,319 | 0,95 | 0,8985 |
| 19 | MAP1LC3B | Microtubule-associated protein 1 light chain 3 beta | NM_001190290 | 21,44 | 0,0185 | 8,37 | 0,0111 |
| 20 | MTOR | Mechanistic target of rapamycin (serine/threonine kinase) | XM_003127584 | 0,76 | 0,6996 | 1,16 | 0,8221 |
| 21 | SQSTM1 | Sequestome-1-like | XM_003123639 | 3,99 | 0,1992 | 0,18 | 0,2985 |
| 22 | TGFB1 | Transforming growth factor, beta 1 | NM_214015 | 0,56 | 0,3277 | 0,18 | 0,0275 |
| 23 | TLR9 | Toll-like receptor 9 | NM_213958 | 2,43 | 0,158 | 0,55 | 0,3275 |
| 24 | UBQLN1 | Ubiquilin 1 | XM_021064658 | 0,46 | 0,3342 | 0,31 | 0,1729 |
| 25 | ULK1 | Unc-51-like kinase 1 (C.elegans) | XM_005670526 | 3,42 | 0,0414 | 1,35 | 0,6935 |
| 26 | ULK2 | Unc-51-like autophagy activating kinase 2 | XM_013981364 | 22,63 | 0,0222 | 2,97 | 0,0167 |
| Oxidative Stress |  |  |  |  |  |  |  |
| 27 | GPX1 | Glutathione peroxidase 1 | NM_214201 | 3,16 | 0,0383 | 2,94 | 0,0496 |
| 28 | GSS | Glutathione synthetase | NM_001244625 | 2,10 | 0,1256 | 1,50 | 0,1256 |

**Table S1**
|  |  |  |  |  |  |  |  |
| --- | --- | --- | --- | --- | --- | --- | --- |
| 29 | Hmox1 | Heme Oxygenase (decycling) 1 | NM_001004027 | 0,73 | 0,4519 | 1,13 | 0,5804 |
| 30 | Nos2 | Nitric oxide synthase 2, inducible | NM_001143690 | 1,48 | 0,4489 | 0,60 | 0,3857 |
| 31 | Sod1 | Nitric oxide synthase 1 (neuronal) | XM_013990334 | 3,36 | 0,0239 | 2,54 | 0,1763 |
| 32 | Sod2 | Nitric oxide synthase 1 (neuronal) | XM_013990334 | 2,74 | 0,0968 | 3,32 | 0,1558 |
| **Apoptosis** |  |  |  |  |  |  |  |
| 33 | AIFM1 | Apoptosis-inducing factor, mitochondrion-associated, 1 | NM_001297632 | 0,96 | 0,9398 | 5,71 | 0,1659 |
| 34 | AIFM3 | Apoptosis-inducing factor, mitochondrion-associated, 3 | XM_013983186 | 1,27 | 0,6638 | 0,60 | 0,4142 |
| 35 | APAF1 | Apoptotic peptidase activating factor 1 | XM_003481742 | 22,85 | 0,1492 | 27,95 | 0,3082 |
| 36 | BAK1 | Bak protein | XM_001928147 | 0,82 | 0,6186 | 0,91 | 0,8137 |
| 37 | BAX | BCL2-associated X protein | XM_003127290 | 16,11 | 0,1216 | 48,06 | 0,2703 |
| 38 | BCL2 | B-cell CLL/lymphoma 2 | XM_003121700 | 5,32 | 0,0209 | 2,30 | 0,08 |
| 39 | CASP3 | Caspase 3, apoptosis-related cysteine peptidase | NM_214131 | 89,44 | 0,0889 | 7,69 | 0,1063 |
| 40 | CASP8 | Caspase 8, apoptosis-related cysteine peptidase | NM_001031779 | 2,28 | 0,292 | 1,79 | 0,4851 |
| 41 | CIDEA | Cell death-inducing DFFA- like effector a | NM_001112696 | 1,07 | 0,9031 | 0,84 | 0,7327 |
| 42 | DIABLO | Diablo, IAP-binding mitochondrial protein | NM_001190216 | 0,49 | 0,0256 | 0,55 | 0,0421 |
| 43 | DFFA | DNA fragmentation factor subunit alpha-like | XM_003127563 | 3,11 | 0,0733 | 1,76 | 0,1588 |
| 44 | FAS | Fas (TNF receptor superfamily, member 6) | NM_213839 | 14,15 | 0,0361 | 0,74 | 0,531 |
| 45 | RIPK1 | Receptor (TNFRSF)-interacting serine-threonine kinase 1 | XM_003128161 | 1,52 | 0,6007 | 0,68 | 0,4966 |
| 46 | TNF | Tumor necrosis factor | NM_214022 | 20,74 | 0,0091 | 14,39 | 0,3567 |
| **Growth factors** |  |  |  |  |  |  |  |
| 47 | FGF2 | Fibroblast growth factor 2 (basic) | XM_013978917 | 0,39 | 0,2403 | 0,52 | 0,3647 |
| 48 | HIF1A | Hypoxia inducible factor 1, alpha subunit (basic helix-loop-helix transcription factor) | NM_001123124 | 9,74 | 0,0432 | 2,93 | 0,42 |
| 49 | IL10 | Interleukin 10 | NM_214041 | 16,16 | 0,1777 | 3,85 | 0,4881 |
| 50 | MMP2 | Matrix metalloproteinase 2 (gelatinase A, 72 kDa gelatinase, 72 kDa type IV collagenase) | NM_214192 | 0,75 | 0,5959 | 1,17 | 0,7673 |
| 51 | MMP9 | Matrix metalloproteinase 9 (gelatinase B, 92 kDa gelatinase, 92 kDa type IV collagenase) | NM_001038004 | 1,01 | 0,977 | 0,54 | 0,1819 |
| 52 | TIMP1 | TIMP metalloproteinase inhibitor 1 | NM_213857 | 1,75 | 0,3061 | 1,11 | 0,8066 |
| 53 | TGFA | Transforming growth factor, alpha | NM_214251 | 1,03 | 0,9594 | 1,22 | 0,7434 |
| 54 | VEGFA | Vascular endothelial growth factor A | NM_214084 | 7,59 | 0,0255 | 1,74 | 0,1304 |
| 55 | NGFR | Nerve growth factor receptor | XM_003131565 | 2,14 | 0,2902 | 0,23 | 0,2853 |
| **Macular degeneration and RPE survival** |  |  |  |  |  |  |  |
| 56 | BEST1 | Bestrophin 1 | XM_005660805 | 5,11 | 0,0045 | 1,37 | 0,4671 |
| 57 | CFH | Complement factor H | NM_214281 | 1,81 | 0,2513 | 5,28 | 0,0263 |
| 58 | ICAM1 | Interacellular adhesion molecule-1 | NM_213816 | 1,31 | 0,7735 | 1,20 | 0,8531 |

**Table S1**
|  |  |  |  |  |  |  |  |
| --- | --- | --- | --- | --- | --- | --- | --- |
| 59 | NOS1 | Nitric oxide synthase 1 (neuronal) | XM_013990334 | 2,67 | 0,0905 | 0,77 | 0,7234 |
| 60 | NOS3 | Nitric oxide synthase 3 (endothelial cell) | NM_214295 | 1,61 | 0,6116 | 5,55 | 0,2175 |
| 61 | RPE65 | Retinal pigment epithelium-specific protein 65 kDa | XM_003127931 | 0,35 | 0,0177 | 1,00 | 0,9974 |
| 62 | TLR3 | Toll-like receptor 3 | NM_001097444 | 3,25 | 0,0299 | 0,36 | 0,0777 |
| 63 | TLR4 | Toll-like receptor 4 | NM_001113039 | 1,87 | 0,1172 | 1,56 | 0,2303 |
| **Inflammatory** |  |  |  |  |  |  |  |
| 64 | CCL2=mcp1 | Chemokine (C-C motif) ligand 2 | NM_214214 | 0,64 | 0,3645 | 1,19 | 0,7675 |
| 65 | CXCL2=SDF1 | Growth-regulated protein homolog gamma | XM_005652553 | 0,21 | 0,1877 | 0,12 | 0,1014 |
| 66 | CXCR4 | Chemokine (C-X-C motif) receptor 4 | NM_213773 | 0,18 | 0,3034 | 1,35 | 0,7967 |
| 67 | IL18 | Interleukin 18 (interferon-gamma-inducing factor) | NM_213997 | 12,41 | 0,0215 | 7,29 | 0,0719 |
| 68 | IL17A | Interleukin 17 A | NM_001005729 | 1,96 | 0,4738 | 0,44 | 0,5468 |
| 69 | IL8 | Interleukin 8 | NM_213867 | 6,80 | 0,0327 | 1,76 | 0,2997 |
| 70 | IL1A | Interleukin 1, alpha | NM_214029 | 0,76 | 0,6787 | 0,06 | 0,0239 |
| 71 | IL1B1 | Interleukin 1, beta | NM_214055 | 1,58 | 0,3767 | 1,14 | 0,7981 |
| 72 | IL6 | Interleukin 6 (interferon, beta 2) | NM_214399 | 5,33 | 0,0159 | 9,99 | 0,0978 |
| 73 | IL12RB1 | Interleukin 12 receptor, beta 1 | NM_001145986 | 5,59 | 0,0194 | 2,51 | 0,0309 |
| 74 | TLR2 | Toll-like receptor 2 | NM_213761 | 2,61 | 0,2055 | 0,85 | 0,8103 |
| 75 | TLR1 | Toll-like receptor 1 | NM_001031775 | 1,65 | 0,411 | 2,79 | 0,4546 |
| **Others** |  |  |  |  |  |  |  |
| 76 | ATF4 | Activating transcription factor 4 (tax-responsive enhancer element B67) | NM_001123078 | 4,23 | 0,2829 | 0,04 | 0,2116 |
| 77 | AQP2 | Aquaporine 2 (collecting duct) | NM_001128476 | 0,57 | 0,3246 | 0,32 | 0,0786 |
| 78 | AQP3 | Aquaporine 3 (Gill blood group) | NM_001110172 | 0,53 | 0,1576 | 0,36 | 0,0371 |
| 79 | AQP1 | Aquaporine 1 (Colton blood group) | NM_214454 | 1,10 | 0,7251 | 1,50 | 0,3442 |
| 80 | EGF | Epidermal growth factor | NM_214020 | 3,26 | 0,1327 | 2,36 | 0,2133 |
| 81 | GDNF | Glial cell derived neurotrophic factor | XM_005672424 | 3,47 | 0,0608 | 1,41 | 0,5638 |
| 82 | GFAP | Glial fibrillary acidic protein | NM_001244397 | 2,35 | 0,1292 | 0,46 | 0,1551 |
| 83 | MMP1 | Matrix metalloproteinase 1 (interstitial collagenase) | NM_001166229 | 0,51 | 0,1615 | 1,27 | 0,7891 |
| 84 | MMP3 | Matrix metalloproteinase 3 (stromelysin 1, progelatinase) | NM_001166308 | 0,10 | 0,0076 | 0,12 | 0,0044 |
| 85 | MITF | Microphthalmia-associated transcription factor | NM_001038001 | 1,99 | 0,3944 | 4,88 | 0,0448 |
| 86 | PTEN | Phosphatase and tensin homolog | NM_001143696 | 9,22 | 0,1284 | 5,57 | 0,2605 |
| 87 | PCNA | Proliferating cell nuclear antigen | NM_001291925 | 2,85 | 0,4034 | 0,56 | 0,4127 |
| 88 | SERPINF1 | Serpin peptidase inhibitor, clade F (alpha-2 antiplasmin, pigment epithelium derived factor), member 1 | NM_001078662 | 1,06 | 0,8347 | 0,64 | 0,1454 |

**Table S1**
|  |  |  |  |  |  |  |  |
| --- | --- | --- | --- | --- | --- | --- | --- |
| 89 | SERPINE1 | Serpin peptidase inhibitor, clade E (nexin, plasminogen activator inhibitor type 1), member 1 | NM_213910 | 5,93 | 0,1141 | 2,69 | 0,3189 |
| 90 | NFE2L2 | Nuclear factor (erythroid-derived 2)-like2 | XM_003133500 | 1,84 | 0,4031 | 1,43 | 0,5989 |
| 91 | TIMP3 | Tissue inhibitor of metalloproteinase- 3 | XM_003126073 | 2,53 | 0,3303 | 0,95 | 0,9642 |

